# An inspiration-off attractor supports the robust and flexible control of breathing

**DOI:** 10.1101/2025.09.23.678177

**Authors:** Nicholas E. Bush, Luiz M. Oliveira, Zachary T. Glovak, Jan-Marino Ramirez

## Abstract

Breathing is a fundamental motor rhythm necessary to sustain life. The rhythm and pattern of breathing arises from the coordination of a bilaterally symmetric, rostro-caudally extended column of heterogeneous neural populations in the medulla called the Ventral Respiratory Column (VRC). By recording from the extent of the VRC using Neuropixels during optogenetic and physiological manipulations, and projecting the population activity into a dynamical latent space, we find that GABAergic lung-stretch feedback circuits promote rhythmic population activity by creating a temporary stable fixed point in the latent state that terminates diaphragmatic contraction. Stimulation of GABAergic circuits either intrinsic to the VRC (VRC^GABA^) or via afferent pathways from GABAergic neurons of the Nucleus of the Solitary Tract (NTS^GABA^) advance or delay breathing when activated with specific respiratory phase through temporary stabilization of this fixed point. Lastly, we show that activation of glutamatergic signaling opposes the effects of inhibitory signaling by destabilizing this expiratory fixed point to promote rapid inspiration. Together, we decompose the functional impact of different respiratory control subpopulations to reveal an integrated network level mechanism for respiratory control *in vivo*.

## Introduction

Rhythmicity is both ubiquitous and varied in nervous systems. Rhythmic activity occurs at multiple mechanistic, temporal, and functional levels; from molecules to brain-wide networks, from milliseconds to months, and from stereotyped motor patterns to cognition^2–8^. Consequently, the fundamental questions of how these rhythms are generated and maintained, and how they functionally contribute to an organism’s fitness remains a subject of intense interest. Of these rhythms, the respiratory rhythm is particularly vital. It sustains physiological function and life^9,10^, responds to homeostatic demands and threats like obstruction or irritants^11–14^, serves as a medium for communication^15,16^, and influences our cognitive and emotional state^17–20^. The respiratory rhythm and pattern is generated in a rostro-caudally extended, bilaterally synchronized column in the ventrolateral medulla called the Ventral Respiratory Column (VRC)^21–25^. Central among the VRC structures is the preBötzinger Complex (preBötC), which is intrinsically rhythmic in slices^26–30^ and exerts exquisite control over breathing rhythm and pattern *in vivo*^31–34^. In addition, the components of the VRC are strongly connected within itself, its contralateral counterpart, and myriad other medullary and extra medullary brain regions^35,36^.

The ability to isolate rhythmogenic networks and preserve rhythmogenesis in the absence of sensory feedback^37^ has led to the concept of central pattern generators as the key driver of rhythmic behaviors. In some reduced preparations, (e.g., the *in vitro* medullary slice preparation of a core respiratory, the preBötzinger Complex (preBötC))^27^, intrinsic cellular excitability and/or recurrent excitatory signaling are sufficient for generating rhythms^28,38–43^. In the intact animal, however, recurrent excitatory, inhibitory, and neuromodulatory circuits composed of diverse cell classes with complex connectivity patterns across anatomically distributed brain regions^36,44–48^ coalesce to generate robust, yet flexible motor outputs. Importantly, fundamental attributes of the respiratory rhythm are vastly different *in vivo* and *in vitro,* emphasizing the potential for fundamentally different effective mechanisms to maintain rhythmic activity. For example, the frequency of the generated rhythm is an order of magnitude slower *in vitro* and neural activity becomes essentially silent immediately after termination of inspiration^49^, which is not the case *in vivo*^50^. This leads to the important and largely unresolved question as to how interactions between core rhythmogenic kernels integrate into distributed networks *in vivo* to maintain and control rhythmic functions^48^.

Sensory feedback via incoming projections is a critical component of the respiratory rhythm *in vivo*. Inhibitory circuits, particularly those that relay mechanoreceptive feedback about lung stretch^51–53^, have been implicated as instrumental in other motor rhythms^54,55^, and as a key fast-feedback mechanism which can both depress and accelerate breathing rates on short time scales (within one breath)^49,56^. However, it remains unknown if inhibitory pathways downstream of lung afferents are indeed capable of bidirectional respiratory control^13,57,58^. Further, little is known about how excitatory and inhibitory signaling affects on-going distributed neural population dynamics in the respiratory brain centers *in vivo*.

Recently, we performed large scale recordings along the extent of the VRC *in vivo* using Neuropixels. By analyzing the simultaneous activity of hundreds of neurons in a low-dimensional, latent projection^59,60^, we showed that the population dynamics of the VRC follow rotational latent trajectories constrained by limit-cycle attractor dynamics^50^. Interestingly, we observed that the trajectories of the neural population activity were characterized by a consistent feature. At the end of each inspiration, the neural state accelerated towards a consistent point in the latent space as if the network state was being pushed into an attractive well. We hypothesize that this acceleration is instrumental in setting the respiratory rate, and is triggered by timed, phasic inhibition. We further hypothesize that inhibitory populations in a restricted part of the Nucleus of the Solitary Tract at the level of the Area Postrema (NTS - second order lung stretch relay neurons, i.e. pump cells^61,62^) could elicit both bidirectional breathing rate modulation and trigger the characteristic acceleration of the latent trajectory.

Here, we test these hypotheses by performing high-density neural recordings with Neuropixels from the extent of the VRC during direct activation of either inhibitory signaling in the contralateral VRC (VRC^GABA^), inhibitory NTS (NTS^GABA^) populations, or the endogenous lung stretch feedback circuit via the physiological lung stretch defense reflex (Hering-Breuer reflex^63^).

We show that precisely timed phasic optogenetic stimulation of VRC^GABA^ or NTS^GABA^ neurons elicited bidirectional control of respiratory rate, confirming the sufficiency of afferent feedback to both depress and accelerate breathing. Further, all three manipulations caused latent trajectories to accelerate to the same fixed point, effectively “short-cutting” the eupneic limit cycle. This fixed point coincided with the neural state at the offset of inspiration. Importantly, activation of the NTS^GABA^ populations exhibits an exquisite sub-phase specificity of effects observed in the pattern of respiratory rate changes, single neuron firing rates, and latent trajectories that contrasted with the activation of VRC^GABA^. These data highlight that activating inhibitory inputs projecting to the VRC is quantitatively and functionally distinct from activating inhibitory neurons within the VRC itself, despite the apparent similarities of their effects on respiration. Importantly, we show that direct activation of VRC inhibition exhibits inspiration promoting effects not present when stimulating the afferent input. That is, local inhibition in the VRC keeps the network state poised on the edge of an inspiration, while incoming inhibition from NTS^GABA^ forces the state away from the expiration/inspiration transition.

We further contrast the differential effects of inhibitory neurons with the results obtained by the direct activation of excitatory, Vglut2^+^ neurons in the VRC (VRC^VGLUT2^). This caused expected amplitude and respiratory rate increases^31,49^ and coincided with a repulsion of latent trajectories from the inspiratory-off fixed point. Subsequently, latent neural trajectories were not short-cut; rather, trajectory speed increased, representing a qualitatively different network mechanism contributing to respiratory rhythmogenesis.

Lastly, we develop a data-driven, generative dynamical model^64,65^ that reveals that inhibitory and excitatory stimulations can be interpreted as uniform vector fields perturbing the on-going eupneic limit cycle. Importantly, these generative models extrapolate to predict experimental data not present in the fitting procedure (e.g., bidirectional respiratory control and reset curves), as well as a relationship with stimulus amplitude described elsewhere^25^. Excitation and inhibition emerge to be diametrically opposed in this dynamic representation, and stabilize or destabilize, respectively, the inspiratory-off fixed point. Consistent with the quantitative difference between direct or indirect inhibition of the VRC, we further distinguish the resultant stimulus vector fields that emerge during VRC^GABA^ and NTS^GABA^ stimulations. This distinction identifies inspiration promoting components of inhibitory signaling within the VRC.

Together, these data support and expand the classical inspiratory off-switch hypothesis^56,66^ with cell-type specific manipulations and neural population recordings at single cell resolution. This hypothesis states that the normal breathing rhythm is governed by a lung-volume set point that triggers a reset of the inspiratory cycle when reached. We show that activation of putative pump cells is sufficient for this behavior and results in reset curves consistent with an inspiratory-off switch. Importantly, we offer a reframing of this hypothesis from the view of dynamical systems. The core rhythm and pattern generating circuitry in the VRC can be represented as a low-dimensional limit cycle. When inspired air generates sufficient lung stretch that is relayed by afferents, the VRC network state is forced toward a more stabilized inspiratory off fixed point. Upon ceasing inspiratory effort, the lung stretch signal diminishes and inspiration promoting mechanisms prevail to initiate the next breath. However, if lung stretch remains high (as it does during triggering of the Hering-Breuer reflex), the inspiratory-off point remains stable, preventing or slowing respiration. This inspiratory-off switch/attractor hypothesis is corroborated by experimental evidence that respiratory rates decrease, and motor amplitudes increase, when vagal afferents are removed^49,67^.

## Results

### Activating inhibitory populations in the Nucleus of the Solitary Tract drives phase-dependent changes in breathing distinct from activation of excitatory and inhibitory neurons in the VRC

To test the effects on the VRC neural dynamics of perturbing constituent components of the brainstem respiratory neural circuitry, we inserted Neuropixel 1.0 probes along the length of the left VRC^50^ of urethane anesthetized mice while optogenetically stimulating inhibitory (Vgat^+^) neurons or excitatory (Vglut2^+^) neurons of the contralateral (right) VRC (VRC^GABA^, VRC^VGLUT2^ respectively, or Vgat^+^ neurons of the Nucleus of the Solitary Tract (putative pump cells, NTS^GABA^). To target VRC^GABA^ neurons, we generated Vgat^Cre^;Ai32 (n=8) or Vglut2^Cre^;Ai32 (n=7) crosses and acutely inserted a 200*μm* optical fiber over VRC (Figure 1A, Supplemental Figure 1). To target NTS^GABA^ neurons, we injected Cre-dependent ChRmine bilaterally into the NTS of Vgat^Cre^ mice (n=9). ChRmine-oScarlet was well expressed in the NTS (Supplemental Figure 1D); some animals were only transfected unilaterally, but similar physiological responses were observed, thus bilateral/unilaterally transfected mice were pooled.

**Figure 1:**
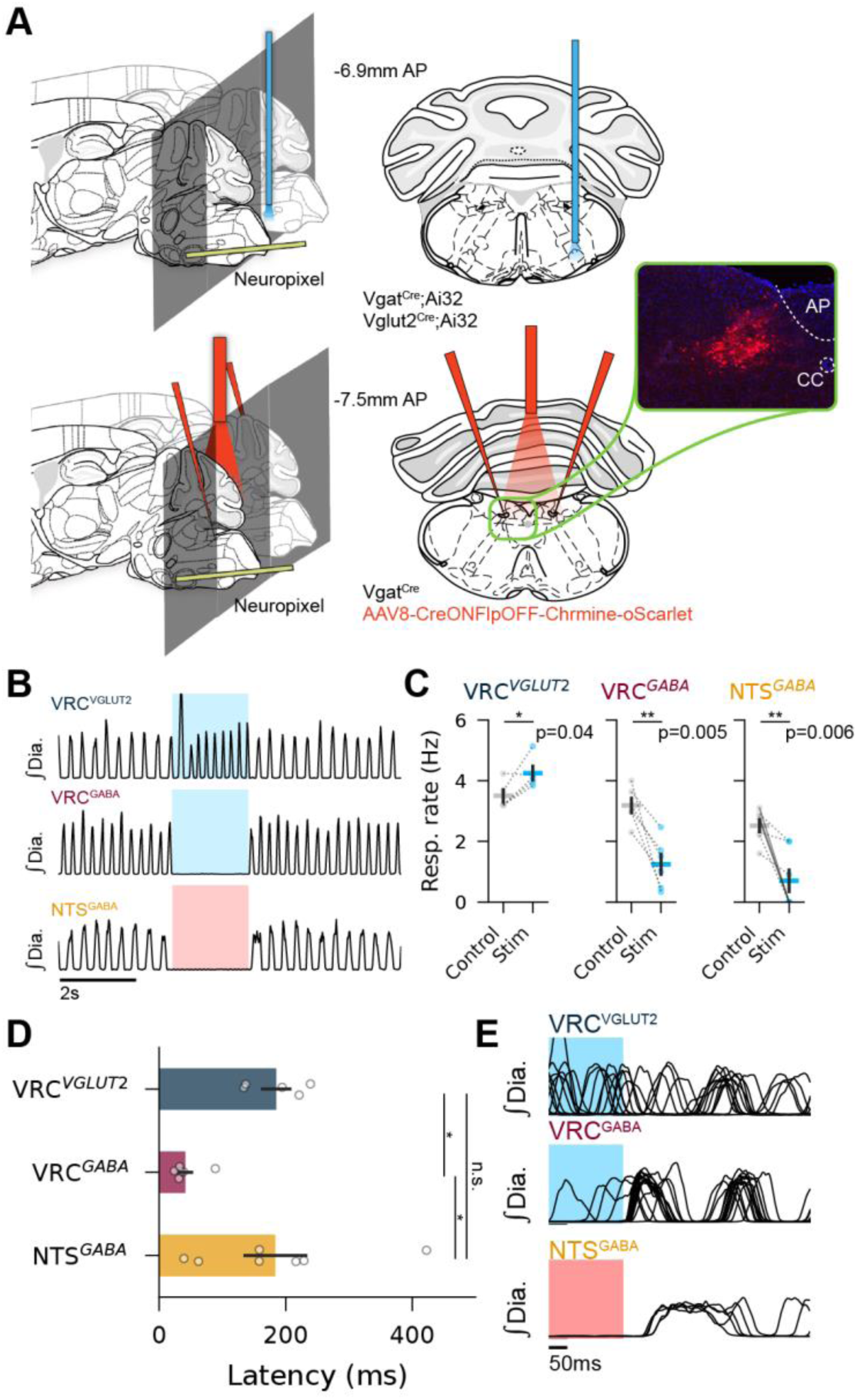
Activation of ventral and dorsal medullary groups differentially affect breathing. (A) Experimental schematic (B) Example integrated diaphragm traces during optogenetic stimulation. Shaded region is laser on. Color indicates wavelength used (blue:473nm, red:635nm). (C) Mean (+/- S.E.M.) respiratory rate during before and during optogenetic stimulation during activation of different populations. Dots are individual animals. Solid lines indicate bilateral transfection/stimulation. (Paired t-test two-sided, VRC^VGLUT2^ n=5, p=0.036; VRC^GABA^ n=6,p=0.004; NTS^GABA^ n=7, p=0.006) (D) Latency (mean +/- S.E.M.) from laser offset to onset of first post-stimulation breath. (Kruskal-Wallis test H=10.2, dof=2,p=0.006; *p<0.05 Holm-Bonferroni corrected) Dots are individual animals. (E) Number of transfected neurons in the NTS as a function of bregma. Black line is mean (shaded region +/- S.E.M.), colored lines are individual animals. Atlas images adapted from^1^

In agreement with previous reports, activating VRC^VGLUT2^ neurons with 2s of constant laser illumination increased respiratory rate, while activating inhibitory neurons in the VRC or the NTS decreased or abolished breathing (VRC^GABA^ or VRC^VGLUT2^) 473nm, 8-10mW; NTS^GABA^ 635nm, 15-20mW)^13,58^ (Figure 1B, C). Activation of NTS^GABA^ slightly decreased, and activation of VRC^VGLUT2^ increased, heart rate (Supplemental Figure 2B). Interestingly, the latency to the first breath after cessation of stimulation was shorter when stimulating VRC^GABA^ populations compared to either VRC^VGLUT2^ or NTS^GABA^ populations (Kruskal-Wallis test p<0.006). (Figure 1D, E).

We next replicate the phase specific effects of stimulating VRC inhibitory or excitatory populations^49^, and tested if stimulation of NTS^GABA^ populations exhibit the effect on breathing as stimulating directly VRC^GABA^ neurons. As expected, activation of VRC^VGLUT2^ neurons during inspiration does not change respiratory rate, but increases diaphragm EMG amplitude; and stimulation during expiration increases respiratory rate (Figure 2A, D; Supplemental Figure 2A). Conversely, stimulation of VRC^GABA^ and NTS^GABA^ neurons during expiration slows respiratory rate, while stimulation during inspiration increases respiratory rate (Figure 2B, C, D). We observed a reduction in diaphragm amplitude during inspiratory stimulation of both populations, while we only observed a reduction in amplitude during expiratory stimulations during activation of NTS^GABA^ populations (Supplemental Figure 2A).

**Figure 2:**
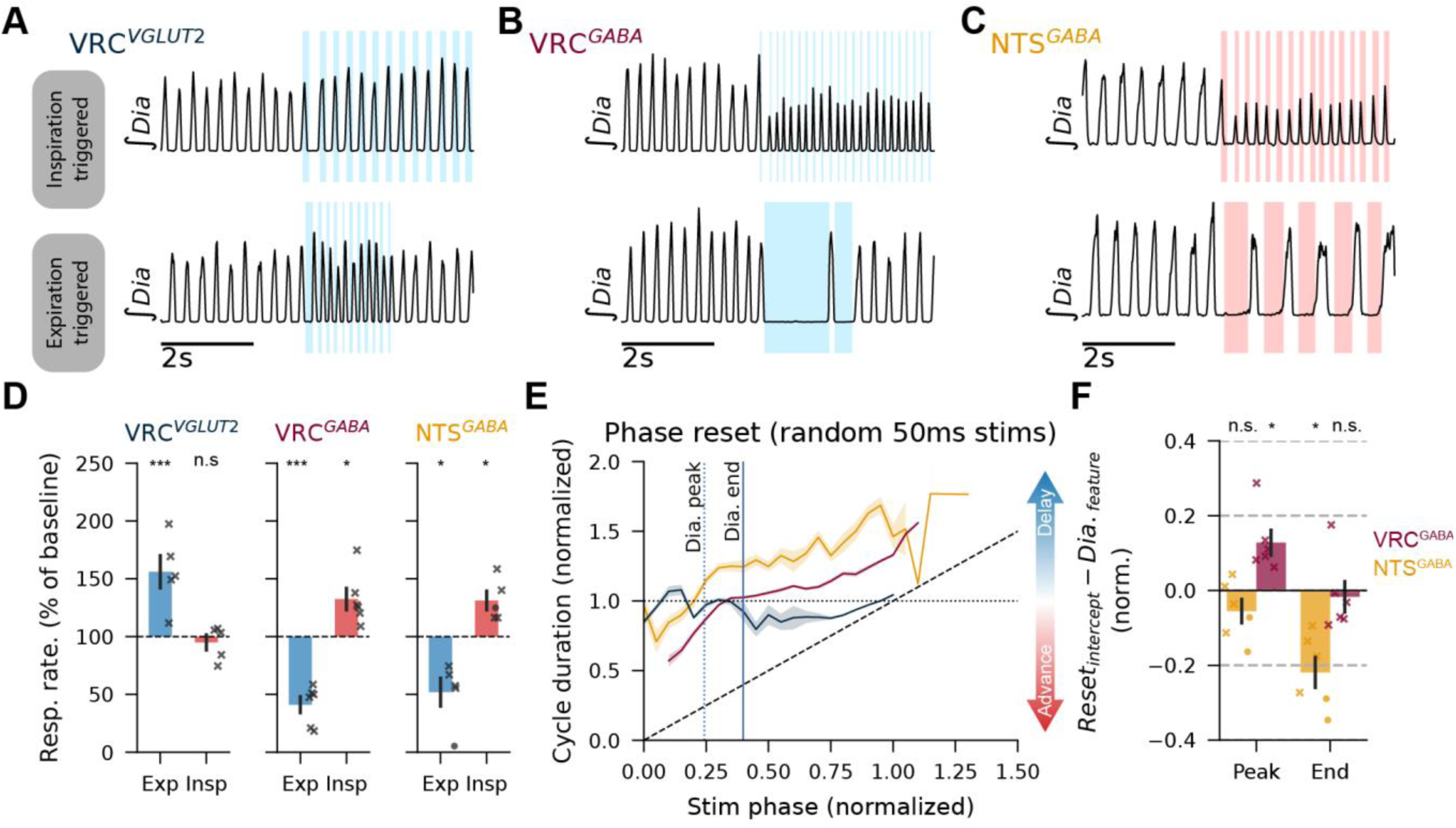
Stimulation of inhibitory NTS neurons exerts phase dependent control of respiratory rate. (A-C) Example integrated diaphragm EMG during laser stimulation of medullary cell types triggered by inspiration (top) or expiration (bottom). Color indicates laser wavelength. (D) Respiratory rate compared to baseline during phase triggered stimulation. (red: inspiration trigger, blue: expiration trigger). (Two-sided one-sample t-test; VRC^VGLUT2^ n=5, p_insp_=0.48, p_exp_=0.016; VRC^GABA^ n=6, p_insp_=0.017, p_exp_=3.7E-4; NTS^GABA^ n=7, p_insp_=0.018, p_exp_=0.017) (E) Phase reset driven by 50ms laser pulses. Color represents stimulated population. Horizontal dashed line represents no change in rhythm, diagonal dashed line indicates theoretical limit on phase advance (i.e., immediate activation of next breath). Vertical lines represent average phase at which diaphragm features occur. (F) Timing of the phase reset curve intercept with 1 (no change in respiratory rate) relative to the peak or end of the diaphragm EMG signal. Color represents stimulated population. (Two-sided one sample t-test; VRC^GABA^ n=6, p_peak_=0.01, p_end_=0.69; NTS^GABA^ n=6, p_peak_=0.14, p_end_=0.003). Marker indicates unilateral (crosses) or bilateral (circles) transfection/stimulation. In all panels, bars/shaded regions are mean +/- S.E.M

Lastly, we presented brief, 50ms pulse laser stimulations (n=75) at random times throughout the phase to observe how momentary activation of these populations perturbs ongoing respiratory rhythms. We compute the reset curves (Figure 2E). Stimulation of VRC^VGLUT2^ neurons advances respiratory phase (curve below 1), specifically when activated during expiratory time (i.e., after the end of the diaphragm EMG activity). Stimulation of VRC^GABA^ advances respiratory phase if stimulating during inspiration and delays respiratory phase if stimulated during expiration. Interestingly, Stimulation of NTS^GABA^ population advances respiratory phase specifically during the first half of the inspiration, i.e., when the diaphragm amplitude is increasing. Otherwise, activation of these populations delays phase. To quantify this difference between activating VRC^GABA^ and NTS^GABA^ populations, we compute the point at which each animal’s phase reset curve crosses one and find that the intercept is at the end of the diaphragm activity for VRC^GABA^ stimulations, but at the peak of the diaphragm for NTS^GABA^ stimulation (Figure 2E, F).

### Activation of NTS^GABA^ populations selectively inhibits inspiratory VRC neural activity

We next asked how activation of these respiratory circuit components drives changes in the activity of recorded VRC populations. These stimulations are targeted to either contralateral (VRC^GABA^ or VRC^VGLUT2^) or input (NTS^GABA^) populations. We analyzed single units recorded from the VRC during the 2s continuous laser stimulations described above. After spike sorting and quality control steps, we categorized single units by their phasic activation patterns as inspiratory, expiratory, or tonic (see methods). Example raster plots showing a subsample of the recorded population before, during, and after optogenetic stimulation show diverse, stimulus specific changes in single unit activity (Figure 3A-C). We compare the firing rate in baseline control conditions against stimulation to see how much each unit is activated/inhibited as a result of stimulating our targeted populations (Figure 3D-F). While there is diversity in firing rate changes in all respiratory populations as a response to all stimulus conditions, we find that activating VRC^GABA^ neurons on average drastically inhibits all neuron types and most neurons in the VRC (Figure 3D, E top). Conversely, activation of VRC^VGLUT2^ increases the firing rate of most neurons on average (Figure 3D, E bottom). Interestingly, activation of NTS^GABA^ populations on average does not change the firing rates of expiratory neurons (Figure 3D, E middle, F top), and only minorly decreases firing of tonic neurons (Figure 3D, E middle, F bottom). Strikingly, these data suggest a functional specificity of activity of NTS^GABA^ populations to selectively decrease inspiratory activity without altering other respiratory population dynamics.

**Figure 3:**
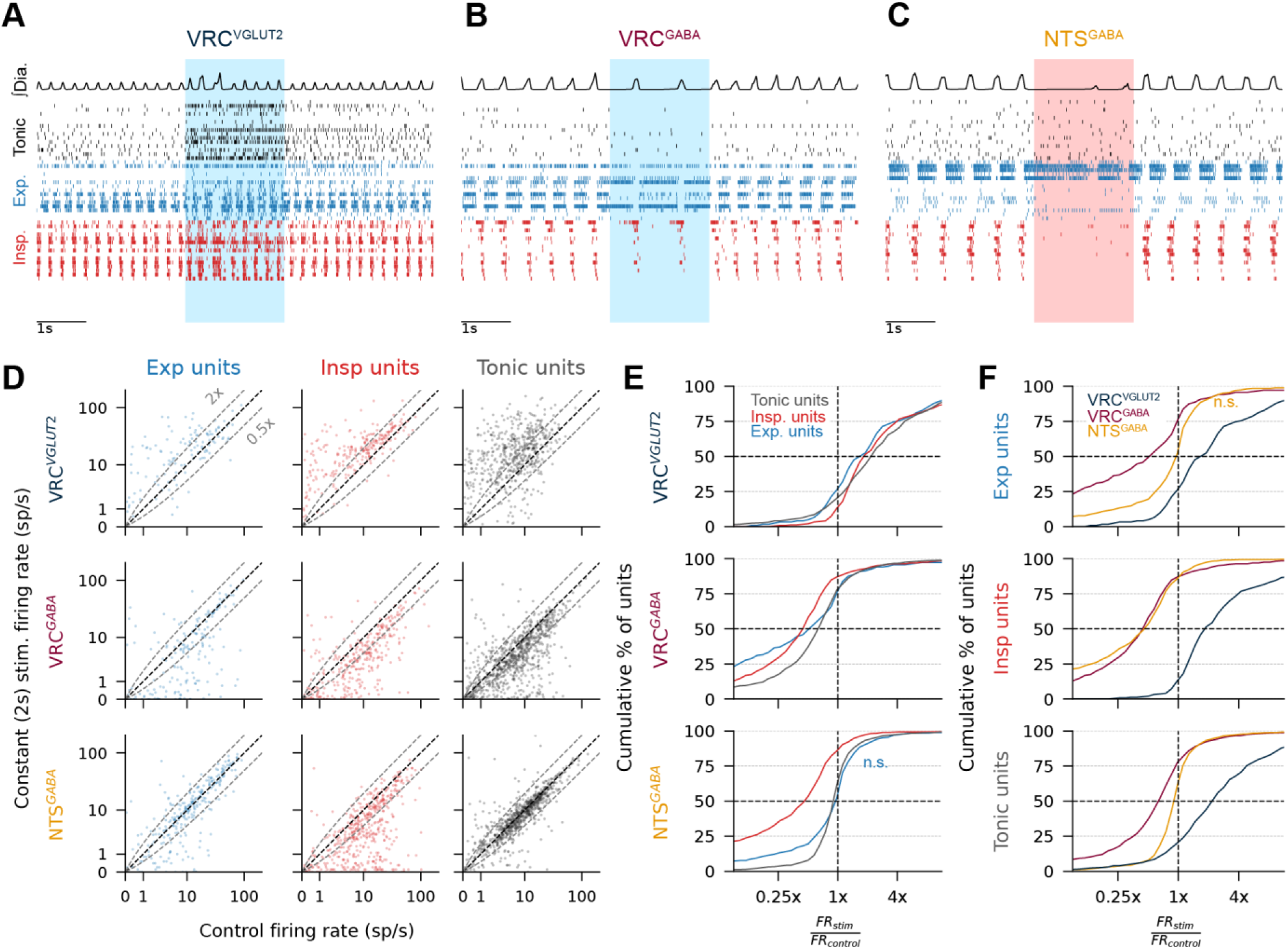
NTS^GABA^ populations selectively inhibit inspiratory neurons. (A-C). Example breathing and activity of example single units of tonic, inspiratory, and expiratory (10 per category, each row is a unit) in the VRC during optogenetic stimulation. (D) Average firing rate (spikes/s) of each unit during baseline control versus during 2s laser stimulations. Each dot is a unit. Note the symmetric logarithmic scale. Black dashed diagonal is unity, and grey dashed lines indicate 2x or 0.5x firing rate during laser stimulus. (E) Cumulative distributions of firing rate ratios (stim vs control). X-axis is log scaled. All comparisons are p<0.05 except expiratory units during NTS^GABA^ stimulation (Wilcoxon signed rank test, control vs stimulus average firing rate). Right shifted curves indicate increases in firing rates during stimulations. (F) same curves as E, but phasic unit types are plotted together across stimulated populations. Color indicates stimulated population.

### Activating inhibitory circuits shortcuts rotational population dynamics towards inspiration off

Recently, we showed that rotational latent dynamics underlie VRC population dynamics^50^. Further, we showed that these rotations targeted an attractive-like region (potential well) that represents the offset of inspiration. Here, we use neuronal subtype specific stimulations to test the hypothesis that activation of inhibitory circuits triggers an acceleration of the latent state towards that inspiration-off target to organize the respiratory rhythm. In addition, we tested a complimentary hypothesis that activation of VRC^VGLUT2^ populations drives the latent state away from the inspiration-off target. Indeed, we observe that activation of both VRC^GABA^ and NTS^GABA^ populations during inspiration drives the network preemptively towards the inspiration-off target, while activation during expiration clamps the network in that potential well (Figure 4A, B middle, right, Supplemental videos 1-2). Activating VRC^VGLUT2^ populations does not alter the rotation but does increase trajectory speed (Figure 4A, B left, Supplemental Figure 3, Supplemental Video 3). NTS^GABA^ stimulation most dramatically reduces trajectory speed during pre-inspiration and inspiration (Supplemental Figure 3C, D), emphasizing the selective inhibition of expiratory dynamics. We next examine the trajectories during and after the randomly timed, 50ms laser pulses. Agreeing with the phase trigged stimulus results, activating both VRC^GABA^ and NTS^GABA^ populations drives the network directly toward the inspiration-off target, while activating VRC^VGLUT2^ populations drives the network along its current trajectory. We quantify the degree of convergence of the trajectories by measuring the dispersion of the two-d latent trajectories 25ms after the offset of the stimulus pulse (i.e., to address the question how similar the network states are shortly after the stimulus, see Methods). Activating VRC^VGLUT2^ populations does not reduce the dispersion compared to randomly selected states during baseline, while activating either inhibitory population does. Similarly, we compute the distance from the post-stimulation state to the inspiration-off point observed during baseline breathing and observe VRC^VGLUT2^ pushes the network away from the inspiration-off state toward inspiration peak, while both inhibitory populations push the network toward inspiration-off (Figure 4E, Supplemental Figure 4A). Lastly, we compute this distance as a function of post stimulus time (Figure 4F), and of latent state dimensionality (Supplemental Figure 4). Interestingly, VRC^GABA^ converges to the inspiration-off target more quickly and diverges more rapidly than the NTS^GABA^ stimulations, suggesting an underlying difference in how these populations affect VRC dynamics.

**Figure 4:**
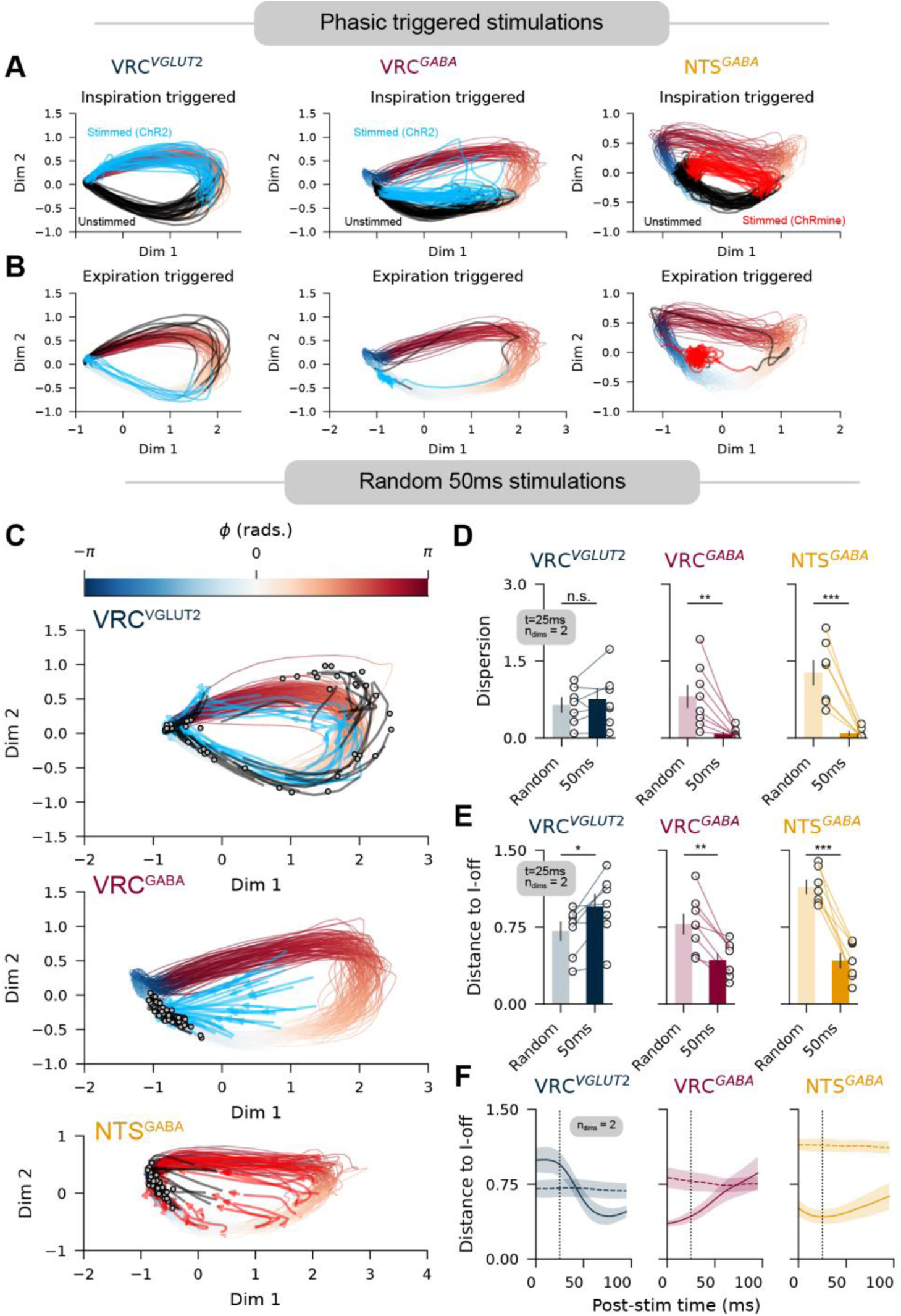
Inhibitory stimulations drive VRC populations dynamics towards inspiration off. (A) Latent dynamic trajectories during inspiration and (B) expiration triggered stimulations of medullary populations for three example recordings. Blue-white-red gradient shows normal latent trajectory (color mapped to respiratory phase), black lines indicated trajectory during unstimulated periods, blue/red lines indicate trajectories during stimulated periods corresponding to laser wavelength used to stimulate respective opsins. One stimulation replicate is shown (inspiration triggered: 15s, expiration triggered: 2s). (C) Randomly timed, 50ms pulse stimulations (75 pulses) of each medullary population. Red-white-blue gradient lines indicate normal trajectory, blue/red lines indicate trajectory during stimulation as in A. Black lines indicated trajectory for 25ms after end of stimulation. Grey dots indicate latent state 25ms after end of stimulation. (D) Dispersion (i.e., trace of the covariance matrix) of two-dimensional latent state a t=25ms post-stimulation, compared to a sample of randomly chosen points in the baseline period. Each dot is a recording. (Paired t-test two-sided, VRC^VGLUT2^ n=7, p=0.24; VRC^GABA^ n=8,**p=0.006; NTS^GABA^ n=7, ***p=8.3E-4). (E) Distance between average two-dimensional latent state at inspiration offset compared to 25ms post-stimulation. Random shuffled control as in D. (Paired t-test two-sided, VRC^VGLUT2^ n=7, p=0.014; VRC^GABA^ n=8,p=0.0014; NTS^GABA^ n=7, p=2.7E-5). (F) Distance to the inspiration off average latent state as in E, for all post-stimulation time delays. Dashed line indicates random shuffled controls. Lines, shaded regions are mean +/- S.E.M.

### Neural population dynamics during lung stretch reflex suggests inspiratory-expiratory competition mechanism

The Hering-Breuer inflation reflex^63,68^ is characterized by the rapid and robust lung-stretch induced apnea, often caused by exogenous increases in positive airway pressure. This reflex critically protects against damage to the lung due to over inflation. The Neuropixel recordings allow us to determine how triggering this reflex altered neural activity in the VRC populations. We present positive airway pressure (15cm H_2_O) briefly by closing an exhaust valve in the airway line, diverting presented airflow into a water column for 2 or 5 seconds (Figure 5A). This causes a decrease or cessation of inspiration, and concomitant changes in VRC neural activity (Figure 5B, C, Supplemental Figure 5C). Inspirations that break through the inflation stimulation are shorter than control (Wilcoxon signed-rank test p=3.0E-5, Figure 5D, E). We compare the breath-cycle averaged firing rates of respiratory neurons between control and Hering-Breuer stimulation. Inspiratory neurons unsurprisingly have shorter durations of activations during stimulation (Figure 5F, G). The reduction in both phase-normalized inspiratory firing rates (Figure 5H) and average firing rate (Figure 5I), is likely due specifically to a reduction in the inspiration time; computing only the firing rate during inspiration reveals an average increase in inspiration-specific firing rate (Supplemental Figure 5E, F). This indicates that inspiratory neurons fire more spikes per inspiration, but over fewer inspirations with shorter inspiratory times. Interestingly, some expiratory neurons lose the gradual decrement over the expiratory phase that leads towards inspiration (Figure 5F-H) and transition to quiescence more rapidly during the transition into inspiration (Figure 5G, H), signifying a more abrupt state transition. The firing rate of expiratory neurons does not change on average, matching what was seen in direct activation of NTS^GABA^ populations. There is no striking anatomical specificity to changes in neural activity (Supplemental Figure 5A-D) suggesting that the Hering-Breuer affect the entire VRC network.

**Figure 5:**
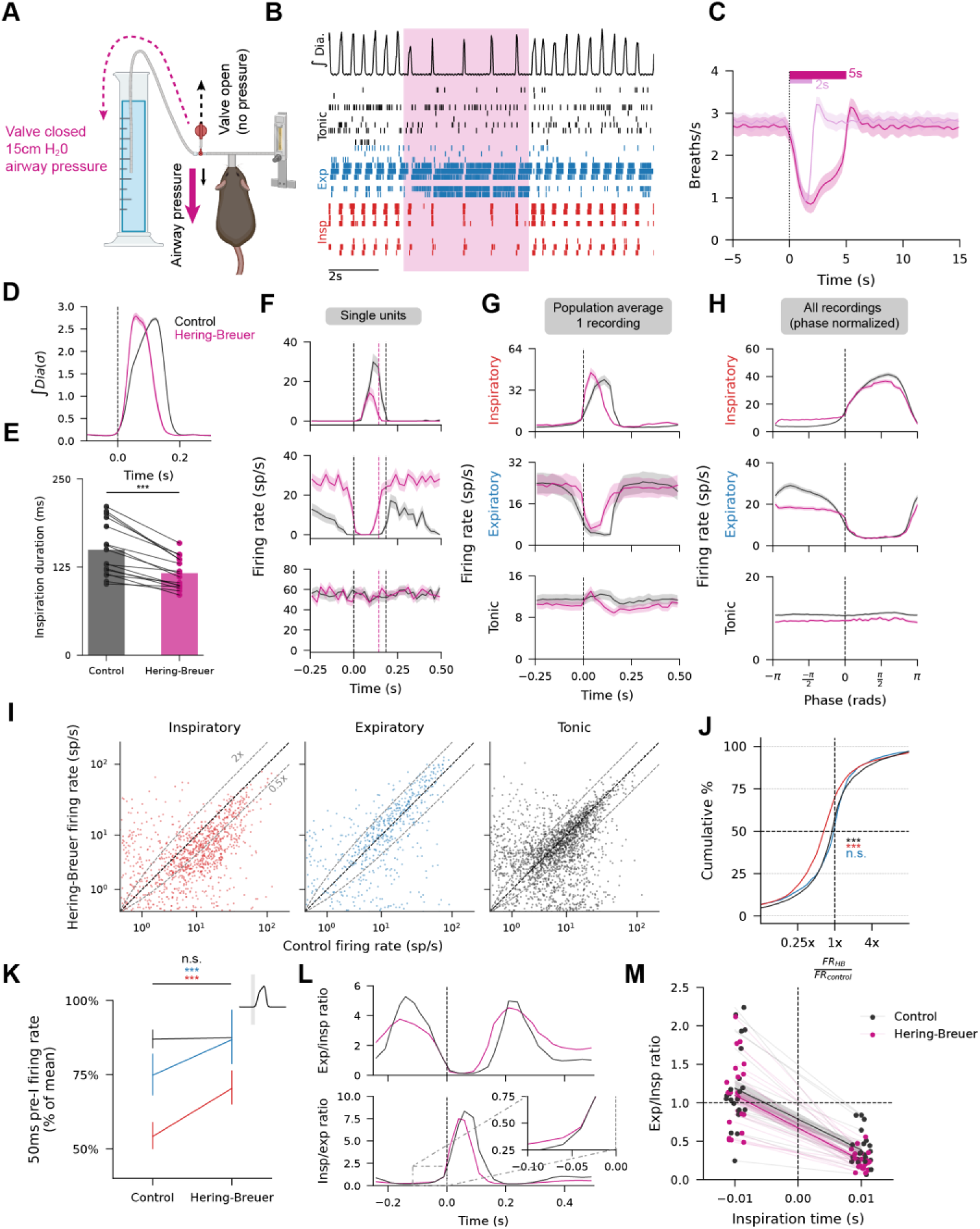
Activation of airway protective Hering-Breuer reflex selectively reduces firing of inspiratory neurons and alters population activity (A) Schematic for triggering positive airway pressure. (B) Example integrated diaphragm EMG and selection of tonic, expiratory, and inspiratory single units during positive airway pressure. (C) Respiratory rate during 2s and 5s positive airway pressure presentation. (n=16). (D) Normalized, breath-triggered average integrated diaphragm EMG for normal breaths (black) and “break-through” breaths during lung inflation (magenta).(E) Average inspiration duration (Wilcoxon signed rank test, n=16,p=3.0E-5). Color as in D. (F) Breath-triggered average firing rates for example single units during baseline (black) and during lung inflation (magenta). (G). As in F, but averaged over all neurons of a given respiratory category in and example recording. (H) Phase normalized, breath triggered average firing rates for all units across all recordings. Color as in G. Shaded region is mean +/-S.E.M.(I) Firing rate of each unit of a given category in baseline control against during Hering-Breuer stimulation (lung inflation). Note the symmetric logarithmic scale. Black dashed diagonal is unity, and grey dashed lines indicate 2x or 0.5x firing rate during lung inflation stimulus. (J) Cumulative distributions of firing rate ratios (stimulation vs control) (Wilcoxon signed rank test firing rate control vs firing rate during lung inflation insp: Wilcoxon=137,470, ***p=6.4E-48, n=1,064; exp: Wilcoxon=79,908, p=0.43, n=576; tonic: Wilcoxon=976,308, ***p=1.0E-12,n=2,177). (K) Average firing rates for given category (indicated by color as in I) of units in the 50ms before the onset of inspiration (indicated in inset) during baseline control and Hering-Breuer stimulation. Firing rates are a normalized percentage of mean firing rate in baseline control period. (Two-sided paired t-test t_insp_=-5.1,***p_insp=4.9E-7;_ t_exp_=-4.9,_***pexp=1.0E-6; ttonic=1.5, ptonic=0.14)_ (L) Breath-triggered ratio of expiratory/inspiratory unit firing rates (top) or *vice versa* (bottom) for all units pooled across recordings during baseline control (black) and lung inflation (magenta). (M) Ratio of firing rates of expiratory neurons to inspiratory neurons 10ms before and after onset of inspiration. Dots/thin lines are individual recordings, thick banded lines are means +/- S.E.M. In all panels, shaded regions are mean +/- 95%CI unless otherwise noted.

To further characterize how lung inflation changes the neuronal network activity during the transition into inspiration, we compute the firing rate of each neuron type (inspiratory, expiratory, and tonic) in the 50ms before the onset of inspiration. Both inspiratory and expiratory neurons show higher firing rates in this period during the lung inflation stimulation compared to control (Figure 5K, two-sided paired t-test p_insp_=4.9E-7 p_exp_=1.0E-6). Lastly, we compare the ratio of inspiratory to expiratory firing rates, pooled across all neurons and recordings. Interestingly, we find that the inspiration on transition occurs when the inspiratory/expiratory firing rate ratio is one in both control and stimulated conditions (Figure 5K, M), irrespective of the observation that the absolute firing rates are higher.

### Lung-inflation induced latent dynamics resemble direct stimulation of inhibitory populations

We next examined the low-dimensional latent state dynamics of the VRC population during Hering-Breuer reflex activation evoked by lung inflation stimuli. Presentation of positive pressure causes that latent state to arrest at a single point in the latent space (Figure 6A, Supplemental Video 4). This arrest is exemplified by a reduction in latent trajectory speed (Figure 6B) that matches well the arrest observed when we stimulated either the VRC^GABA^ or NTS^GABA^ populations directly. (Figure 6C, D, also compare to Figure 4B). In addition, trajectory speed during inspiration onset is higher during Hering-Breuer break through breaths (Supplemental Figure 5G). The lung inflation stimulation converges the latent state toward the inspiration-off attractive target (Figure 6E, F) as in the stimulation of inhibitory populations. Lastly, we asked if the single neuron activity observed during lung-inflation stimulations resembled the single neuron activity observed during the 2s optogenetic stimuli applied previously (Figure 6G). We compute the difference and Pearson correlation of the mean square-root transformed firing rates between the two stimulation paradigms. The difference indicates if the average firing rates are similar across conditions, and the Pearson correlation indicates if individual units are modulated similarly across the different conditions. While VRC^VGLUT2^ and VRC^GABA^ showed higher and lower average firing rates than the lung inflation stimulation, respectively for all respiratory neuron types, NTS^GABA^ stimulation matched the average firing rates well (Figure 6H). Interestingly, the correlation of the inspiratory neurons during NTS^GABA^ stimulation was strikingly lower than expiratory or tonic populations, while that of the inspiratory neurons during VRC^GABA^ stimulation was high, suggesting incomplete mimicry of the physiological stimulus with this circuit specific activation, despite a strong match in the physiology.

**Figure 6:**
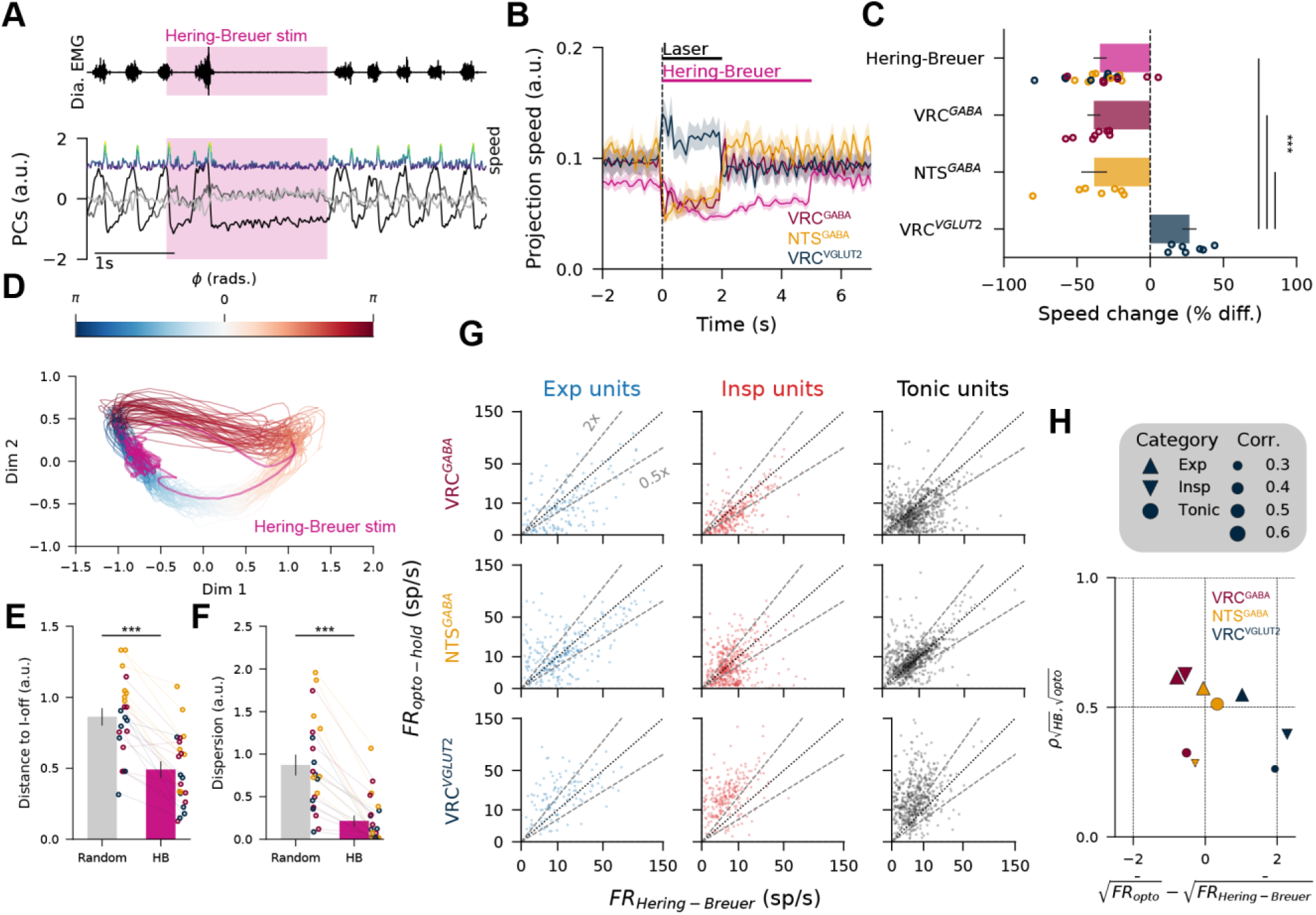
Lung inflation drives latent dynamics towards inspiration off attractor similar to activation of inhibitory circuits. (A) Example diaphragm EMG (top), first 3 PCs (grayscale), and latent trajectory speed (blue-yellow gradient, scale arbitrary, not shared with PCs) during a 2s lung inflation. (B) Trajectory speed of leading 3 dimensions for all recordings during 2s laser stimulations (color indicates targeted population), or 5s Hering-Breuer stimulation (magenta). Shaded regions are mean +/- S.E.M. (C). Percent change in trajectory speed in the period (1,2)s during stimulation compared to the period (-2,-1)s before onset of stimulation. One-way ANOVA (F=25.5, DF=3,DF=40,p=2.2E-9; Tukey-HSD post-hoc test ***p<0.001). (D) Example two-dimensional latent trajectories during baseline (red-white-blue gradient lines colored by phase) and during one 5s lung inflation stimulation (magenta). (E) Distance between average two-dimensional latent state at inspiration offset compared to random shuffled points (Random) or 1s after lung inflation onset (HB). Dots, thin lines are recordings. Color indicates optogenetically targeted population in other experiments. Linear mixed effects model DF1=2,DF2=19; F_population_= 9.14, p=1.6E-3; F_HBstim_=56.33,***p=4.3E-7; F_interaction_=1.4,p=0.27. (F) As in E, but comparing the dispersion (i.e., trace of the covariance matrix) of two-dimensional latent state a 1s after lung inflation onset. Linear mixed effects model DF1=2,DF2=19; F_population_= 2.14, p=0.145; F_HBstim_=56.2,***p=4.3E-7; F_interaction_=1.98,p=0.15). (G) Firing rate of individual units during lung inflation stimulation (x-axis) and during 2s optogenetic hold stimulation (y-axis), stratified by stimulated population and respiratory neuron type. Note the square root scaled axis. Black dashed diagonal is unity, and grey dashed lines indicate 2x or 0.5x firing rate during optogenetic stimulation compared to lung inflation stimulation. (H) Comparison of firing rates of subpopulations during different stimulus modalities. X-axis shows difference of the means of square root firing rates between optogenetic and lung inflation stimuli. Zero indicates similar firing rates in both conditions; positive values indicate higher firing rates during opto-genetic stimulus. Y-axis and marker size show correlation (*ρ*) between the firing rates of the two modalities. Color shows optogenetically targeted population, marker shape indicates respiratory category.

### Activation of brainstem subpopulations are transient forces applied to ongoing dynamic landscapes

We previously reported that explicitly incorporating dynamics into our latent state models reveals a limit cycle underlying VRC population dynamics that is highly consistent across animals^50^. This limit cycle was well recovered with a recurrent switching linear dynamical system (rSLDS)^64^ with two partitions of the latent state (that correspond to inspiration and expiration) and two continuous latent dimensions^50^. Analyses of the latent state above suggest that inhibitory stimulations push the network state back down to the inspiratory off attractor. We test that hypothesis explicitly by incorporating a dependence of the two-dimensional latent state *x*_*t*_ on an external input *u*_*t*_ (see Methods), where *u*_*t*_ is the presence of the laser pulse. We fit the latent state to the baseline (no stimulus) period and periods of stimulation with short (10ms, 50ms) laser pulses. This estimates the latent dynamics, from which we can predict the integrated diaphragmatic activity as a function of that latent state (Figure 7A, B). This fitting also estimates the imposed effect of the optogenetic stimulation on the dynamics (Figure 7B, C, Supplemental Videos 5,6). The learned stimulus effect manifests as two uniform vector fields partitioned by the same partitions learned by the dynamics (i.e., respiratory phase). By increasing the stimulus effect post-hoc, we can elicit qualitative changes in the limit cycle such that the limit cycle disappears (Figure 7B). We first examine the learned stimulus fields. By computing the norm of the stimulus vectors, we find that the VRC^VGLUT2^ stimulus has a stronger effect during expiration, VRC^GABA^ has a stronger effect during inspiration, and NTS^GABA^ has a similar strength during both states (Figure 7D). We next compare the learned inspiratory stimulus direction with the slow eigenvector of the expiratory dynamics. The slow expiratory eigenvector resembles the direction through which the latent state evolves as it transitions into inspiration (Figure 7C). The cosine similarity of the inspiratory stimulus field with the expiratory slow dynamics is large and positive for VRC^VGLUT2^ stimulation, indicating the stimulus pushes the network along the existing dynamics (Figure 7E, Supplemental Figure 6). The stimulus opposes (i.e., large and negative) the normal dynamics during VRC^GABA^ stimulation, and is more diffuse compared to NTS^GABA^ stimulations. These results are consistent with the experimental observations in Figure 4.

**Figure 7:**
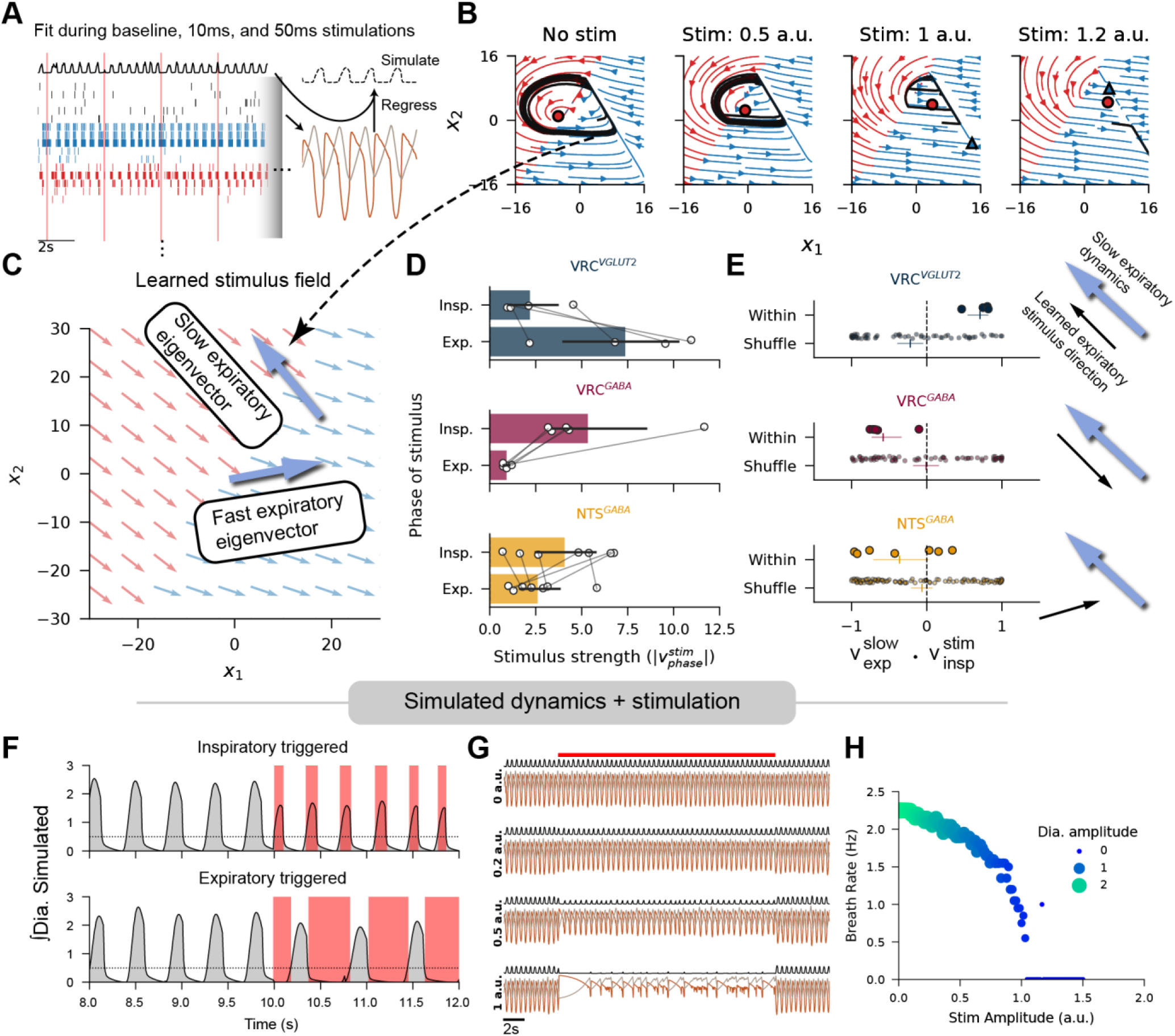
Activation of subpopulations drive generalizable changes to latent dynamics. (A) Example fitting of piecewise latent dynamical system (rSLDS). Example rasters/integrated diaphragm as in Figure 3A. rSLDS model is fit only to spiking observations during baseline, 10ms, and 50ms laser pulses. Fitted latent dynamics are regressed against observed integrated diaphragm to map simulated diaphragm from two-dimensional latent state. (B) Simulated latent dynamics from data in A, varying strength of simulated stimulus. Red and blue streamlines are underlying piecewise dynamical system, black lines are example latent evolutions for 5s. Red, blue markers are fixed points of inspiratory/expiratory partitions of latent state, respectively. Circle is stable node, triangle is saddle. (C) Red and blue arrows represent the learned effect of optogenetic stimulation on the latent dynamics, as they affect the inspiratory/expiratory partitions respectively. Light blue arrows show the two eigenvectors of the expiratory partition in B. (D) Norm of the stimulus (i.e., stimulus strength) as it affects the dynamics of inspiratory or expiratory partitions, stratified by stimulated populations. Each connected pair of dots is one recording. (E) Cosine similarity (i.e. dot product of normed vectors) comparing the direction of the slow expiratory eigenvector with the direction of the inspiratory stimulus field. Each dot is a recording. Shuffle compares learned stimulus with eigenvectors from other recordings. (F) Example generalization of learned stimulus effect to recapitulate phase-dependent effects on respiratory rate in a simulated NTS^GABA^ recording, simulated entirely from learned dynamics. Gray trace is simulated diaphragm, shaded red regions are triggered stimulations(G) Example generalization of varying stimulus strength in a simulated NTS^GABA^ recording, gradually decreasing and abolishing respiration. Brown, grey traces are simulated latent, black trace is simulated diaphragm. Red line is duration of applied stimulation. (H) Quantification of results in G with high-density parameter sweep. Dot size and color indicate amplitude of simulated diaphragm during inspirations. In all panels, bars are mean +/- 95%CI.

We next asked if applying this learned stimulus field could generalize to other types of stimulus patterns that were not part of the fitting procedure. We applied phasic simulated NTS^GABA^ stimulations and found that indeed, inspiratory triggered stimulations increase simulated respiratory rate, and *vice versa* (Figure 7F). We then simulated long stimulus holds for a range of amplitudes (Figure 7G, H) and found that both simulated respiratory rate and amplitude decrease gradually as a function of stimulus strength until it is abolished. This amplitude sweep was not collected here for direct comparison, but matches published data in^25^.

### Dynamic motifs generalize across stimulus types to predict neural activity and physiological behavior

Strikingly, we observed that the learned stimulus fields are qualitatively similar within a given experimental stimulus (e.g., VRC^GABA^ stimulus fields oppose the slow expiratory dynamics (Figure 7E). We asked if we could apply arbitrary stimulus fields that are defined based only on the observed dynamics without fitting the stimulus directly. If so, we could, for example, artificially apply stimulation of all three experimental populations in the same experimental recording. We generated three prototypical stimulus fields: VRC^VGLUT2^-like VRC^GABA^-like, and NTS^GABA^-like (Figure 8A). These were defined as being: inline with slow expiratory eigenvector only during expiration, opposed to the expiratory eigenvector only during inspiration, and opposed to the expiratory eigenvector over the whole space, respectively (see Methods). Thus, we fit the dynamics only to the baseline recording, computed and applied these stimulus fields for each recording, and simulated diaphragmatic activity through our regression model. Strikingly, we could well recreate many of the observed experimental phenomena: reset curves as created with 50ms stimulations (Figure 8AB), phase dependent modulation (Figure 8C, D, Supplemental Figure 6C, E), rebound latency (Supplemental Figure 6D), and stimulus amplitude dependence (Figure 8E, F). A notable difference is that large stimulus simulation of VRC^VGLUT2^-like fields caused a reduction or cessation of respiratory rate. We expect his was due to a limitation of the piecewise linear model; the state would become restricted to the inspiratory region of the latent space, which was often a stable spiral and would subsequently fail to reach the expiratory region. Together these results present a predictive, network-level control mechanism for respiratory neural dynamics by which subpopulations exert specific influences on the dynamic landscape to reshape respiratory rhythms.

**Figure 8:**
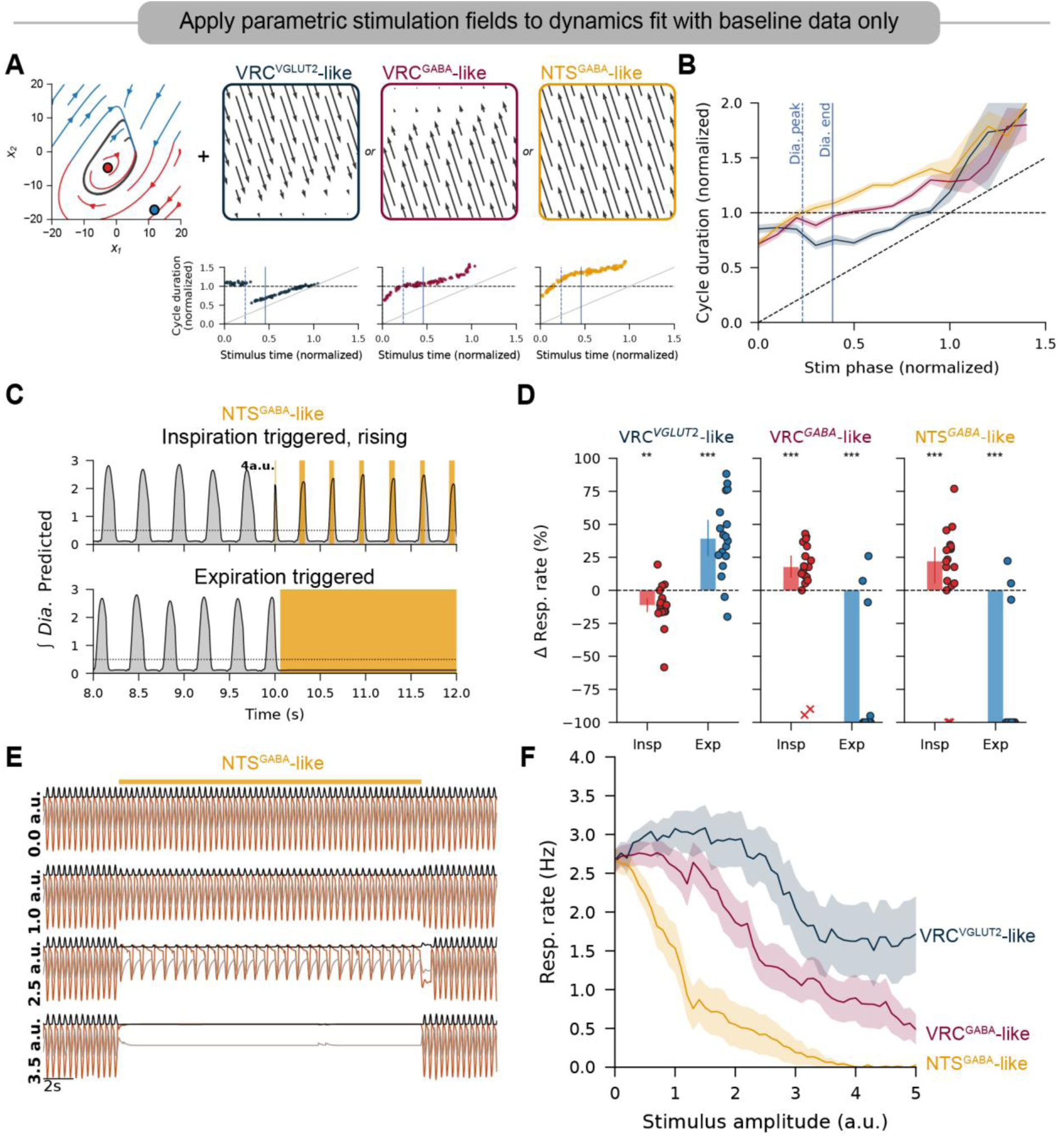
Experimental stimulation of specific subpopulations is recapitulated by parametric, unlearned perturbations of underlying dynamical systems. Simulated stimulations are indicated by green. (A) Latent dynamics are fit without inclusion of optogenetic stimulation. (Left) Dynamics for an example recording (red/blue streamlines, fixed points as in 7B) and (right) the three candidate stimulus fields (black arrows). Direction of stimulus field is always aligned to slow expiratory eigenvector; same direction for VRC^VGLUT2^-like, opposite direction for inhibitory-like. (bottom) Simulated phase reset curves for application of these stimulus fields (50ms stimulation). Reset curves as in Figure 2E. Stimulus amplitude is set to 4. (B) Average reset curves applying the three stimulus fields in A to all recordings. Shaded region is mean +/- 95%CI. All simulated fields generalize to all recordings, regardless of animal genotype recorded; n=19 per group. (C) Example simulated phasic stimulation using an NTS^GABA^-like (i.e., uniform) stimulus field. (D) Change in simulated respiration rate for the three stimulus fields applied during inspiration or expiration (each marker is a recording, crosses are omitted from statistics as outliers, two outlier points of Δ Resp.Rate >400% are omitted). Stimulus amplitude is set at 4. (One sample Wilcoxon signed rank test **p<0.01,***p<0.001, see source data for p-values). Bars are median +/- 95%CI (E) Example varying strength of NTS^GABA^-like stimulus for held period. Brown, grey traces are simulated latent, black trace is simulated diaphragm. Top gray line is duration of applied stimulation. (F) Average simulated respiratory rate as a function of supplied stimulus amplitude for the three stimulus types (color as in B, n= 19 per stimulus application). Shaded regions are mean +/-S.E.M.

## Methods

### Animals

All procedures were approved by the Seattle Children’s Research Institute Institutional Animal Care and Use Committee. Male and female mice aged 9-24 weeks were used. For targeting of VRC^VGLUT2^ and VRC^GABA^ populations, homozygous Vglut2^Cre^ (Jax #028863) and Vgat^Cre^ (Jax# 016962) were crossed with homozygous Ai32 mice (Jax# 012569) allowing Cre-dependent expression of a channelrhodopsin-2/EYFP fusion (ChR2(H134R)/EYFP).

### Surgery

To target NTS^GABA^ populations, we injected AAV8-nEF-Con/Foff 2.0-ChRmine-oScarlet (30 nL; 137161-AAV8) into the NTS at the level of the Area Postrema (7.56 mm caudal from Bregma, ±0.5 mm lateral from the midline and 3.9 mm below the dura mater) of Vgat^Cre^ (Jax# 016962) mice. Anesthesia was induced with 3% isoflurane and maintained at 1.5%. Body temperature was maintained at 37°C with a recirculating heated waterbed (Adroit medical). Peri-operative Ketoprofen (5 mg/kg sc.) was administered. Scalp fur was shaved and the scalp was sterilized with isopropyl alcohol and betadine prior to incision. The skull was cleared and a midline craniotomy extending over the left and right NTS was drilled (± 0.7 mm from the midline). Virus was loaded into a glass pipette attached to a Nanoject 2 (Drummond Scientific), and the pipette was targeted to the left NTS. After cranial injection, the pipette remained in place for ∼5 minutes prior to removal of the pipette and targeting of the right NTS. Skin was then sutured over the craniotomies and mice were allowed to recover. Ketoprofen (5 mg/kg sc.) was administered for two consecutive days after the surgery. Virus was allowed to transfect for 10-28 days prior to recordings.

### *in vivo* Neuropixel recordings

Recording preparations are modified from^50^, see that reference for further details. Anesthesia was induced at 3% isoflurane and urethane (1.5 g/kg) was administered intraperitoneally (IP) before removing animals from isoflurane. Animals were placed on a feedback controlled heating bed (FHC 40-90-8D) to maintain body temperature at 37°C throughout the experiment. Two stainless steel EMG wires (AM systems 0.005 inch diameter) were inserted into the right diaphragm, and two were inserted to record electrocardiogram (one lead over the heart and one in the left intraperitoneal space). EMG wires were secured in the skin with cyanoacrylate (Krazy Glue). EMG signal was amplified (10,000x), hardware filtered (100-5,000 Hz, AM-Systems 1700), and digitized at 10Khz (NI PXIe-8381). Diaphragm EMG was hardware rectified and integrated (custom) to be used for closed-loop online triggering of optical stimulation. (N.B. This rectified and integrated signal was not used in downstream analyses – see Preprocessing). Scalp fur was removed with depilatory cream (Nair). The mouse was then placed in a stereotaxic frame (Stoelting), affixed with earbars, and the skull was leveled. The scalp and skin overlying the rostral neck were removed, and neck muscles were dissected to allow visualization of the foramen magnum and the first cervical vertebra. A stainless steel headplate (custom) was cemented to the skull with UV cure cement (Pulpdent Embrace Wetbond). A small craniotomy was opened over the right cortex, and a bare silver ground wire 0.005in (AM systems) soldered to a gold pin (AM systems) was inserted overlying the dura and secured with cyanoacrylate (Krazy Glue) as a ground. In experiments in VRC^GABA^ or VRC^VGLUT2^ stimulations, a ∼0.5mm craniotomy was drilled over the right VRC (1.25 mm lateral, 6.84mm caudal from bregma) and overlying dura was carefully removed. The animal was then transferred to a custom head-fixing apparatus. A 3D printed nose cone was placed over the animal’s snout, and 100% O_2_ was supplied. If labored breathing was observed, atropine (0.1mg/kg) was administered to alleviate mucous buildup. The dura overlying the left caudal brainstem surface was cut, and the brainstem surface was kept hydrated with PBS. For recordings in VRC^GABA^ or VRC^VGLUT2^ experiments, the optical fiber (200 *μ*m diameter, 0.22NA ThorLabs) was calibrated and then targeted above the right VRC (1.25mm lateral, 6.84mm caudal from bregma, 5.3mm ventral from bregma). In NTS^GABA^ experiments with animals expressing virally mediated, red-shifted ChRmine opsins bilaterally in the NTS, the optic fiber (400*μ*m diameter, 0.22NA, Doric) was placed on the surface of the skull overlying the NTS (midline, 7.2mm caudal from bregma). Neuropixels 1.0 probes coated in DiI (Thermofisher V22885) were then targeted to the left VRC (1.25 mm lateral, 4.8mm ventral from the intersection of the parietal and occipital bones at midline (see ^50^ for details). The probe was then inserted ∼3.5mm rostrally from the caudal brainstem surface at 10*μ*m/s (Scientifica PatchStar). The probe was then retracted 100*μ*m to reduce tissue distortion and allowed to settle for 15 minutes before recording. 10-15 minutes of baseline neural and respiratory activity were recorded before optical stimulation protocols.

### Stimulation Protocols

All experimental hardware was controlled with custom python and Arduino software (https://github.com/nbush257/pyExpControl). Red (635nm) or blue (473nm) lasers (Cobalt MLD 635/473) were driven with an analog voltage (0-1v) calibrated to output either: 635nm light 20mW at the tip outside the skull (400*μ*m diameter fiber) or 473nm light 8-10mW at the tip ∼300*μ*m above the VRC (200*μ*m fiber). A 2 millisecond sigmoidal ramp was added to the start and end of each pulse to minimize light artifacts on neural recordings. The following stimulus paradigms were applied:1) Hold stimulus: 5-15 repetitions of 2s illumination interleaved with 20s inter-trial interval. 2) Pulse stimulus: 50 repetitions of 10ms and 50 repetitions of 50ms pulses with 3s inter-trial interval. 3) 10 Repetitions of 10 seconds long inspiratory triggered stimulations. 4) 10 Repetitions of 2-4 seconds long expiratory triggered stimulations.

Phasic (i.e., inspiratory and expiratory) triggered stimulations were performed by applying a hysteresis trigger to the hardware integrated diaphragm. During inspiratory stimulation, a positive threshold crossing turned on the laser and a negative crossing of the signal at 90% of the rising threshold turned off the laser. Expiratory stimulations used the same logic and thresholds, only the laser control was inverted (i.e., laser off when integrated diaphragm was above threshold, laser on when integrated diaphragm was below 90% threshold). Prior to recordings, the experimenter set the threshold manually with the light path interrupted (to not alter respiratory behavior which could alter the threshold choice) such that the threshold was as low as possible while maintaining phasic activation and avoiding constant illumination. To perform positive pressure experiments triggering the Hering-Breuer reflex, nose cone gas outlet was connected to a T-junction where one line led into a 15cm water column and the other to a constitutively open solenoid valve. To apply positive pressure, the solenoid valve was closed, forcing exhaust gas into the water column.

### Histology

After recording, the animal was euthanized, perfused with phosphate-buffered saline followed by 4% PFA, and the brain was removed. The brain was then fixed overnight in 4% PFA, in 15% sucrose for 24 h and finally 30% sucrose. The brain was embedded in embedding medium (NEG-50) and frozen at −80 °C. Then 25-μm sagittal or coronal sections were collected and imaged (Olympus BX61VS) at 4x-10x magnification. Recording location was identified by the DiI fluorescent track and optical fiber placement confirmed by tissue damage. To minimize potential bias in the quantification, all photomicrography and cell counting were performed by a single researcher blinded to the experimental conditions. Cell counts were obtained using ImageJ software (version 1.41; National Institutes of Health). For each mouse, one-in-two coronal series of 25-µm brain sections was analyzed, resulting in sections spaced 50 µm apart. First, section alignment across brains was performed relative to a reference section of the nucleus of the solitary tract (NTS), which is anatomically bordered dorsally by the area postrema, laterally by the spinal trigeminal nucleus, and ventrally by the dorsal motor nucleus of the vagus (DMV). For sagittal sections, all brain slices were analyzed to identify the Neuropixel probe trajectory and optic fiber placement. Sections were mounted onto slides in phosphate buffer (PB) solution in rostrocaudal sequential order. After drying, they were cover slipped with DAPI-Fluoromount (0100-20; SouthernBiotech) and sealed with nail polish. The neuroanatomical nomenclature used in the experiments followed the atlas of Paxinos and Franklin (2012).

### Preprocessing

Preprocessing of physiological and neural data was performed with custom python software available at https://github.com/nbush257/cibrrig. After recording, raw diaphragm EMG was rectified, median filtered, and EKG artifact was removed using custom removal methods. Breaths were then detected using scipy.find_peaks from the resulting timeseries. Neural data were preprocessed using spikeinterface^69^. Briefly, neural traces were destriped: traces were highpass filtered (250Hz), phase shifted, noise channels were removed, spatial high-pass filtered (local median subtraction), and optogenetic-induced artifacts were removed. Recordings were then motion corrected and spike sorted with Kilosort 4^70^. Only units that passed the following quality control metrics were marked as acceptable and included in downstream analyses (noise cutoff < 0.1, sliding refractory period < 0.1, median amplitude > 40.0 *μ*V, number of spikes > 500, presence ratio>0.8; for details see https://spikeinterface.readthedocs.io/en/stable/modules/qualitymetrics.html). For ease of reuse, extracted data were saved according to International Brain Lab Open Neurophysiology Environment conventions (ONE https://github.com/int-brain-lab/ONE).

### Physiology analysis

Respiratory rates were computed during the stimulation period and compared to the equivalent length control period immediately preceding each stimulation. Relative changes in respiratory rate were computed as 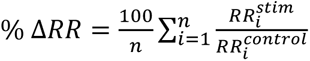 for all *i* stimulations. Similarly for diaphragm amplitude. Reset curves were created by computing the time from each 50ms pulse stimulus to the previous *t_pre_* and the next breath *t_next_* onset, and the average respiratory period 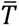. Then we compute normalized values: 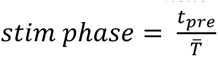 and 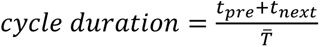. We quantify the intersection (*reset_intercept_*) of the phase reset curve with the *y* = 1 line by first smoothing the reset curve with a second order Savitsky – Golay filter with a window size of 21, then finding the normalized stimulus phase at which that smooth reset curve crosses *y* = 1. We then subtract the normalized phase time to the diaphragm peak, and the diaphragm offset (end). We compute respiratory phase as in^50^ such that respiratory phase *ϕ* is defined on [−*π*, *π*]; *ϕ* = 0 is defined as the onset of an inspiration (i.e., diaphragm activation), and *ϕ* = *π* is the offset of an inspiration (diaphragm inactivation). We set the sample immediately following *ϕ* = *π* to −*π*. Lastly, we linearly interpolate between [−*π*, 0) as expiration, and [0, *π*]as inspiration.

### Single unit analyses

We restrict all analyses to units that passed quality control (see above Preprocessing). The response of each neuron as a function of phase is computed 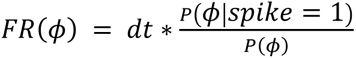 where *dt* is the time step of the *ϕ* signal (1ms) and *ϕ* is binned at a resolution 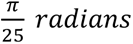. of We define the respiratory modulation and preferred respiratory phase as the vector mean of the phasic histogram from^71^ where the normalized length of the vector is the respiratory modulation ∈ [0,1] and the angle is the preferred phase ∈ [−*π*, *π*]. Units with a respiratory modulation <0.2 were classified as tonic, all others as phasic. Phasic units were classified as inspiratory if the preferred phase was ≥ 0, otherwise expiratory. To compare firing rates between the two stimulus conditions (Hering-Breuer (HB) and opto-hold), we first take the square root transform of the mean firing rate of each unit in the stimulus period to transform the firing rate distributions before computing the correlation *ρ*. We also compute the difference in the mean square root firing rates: 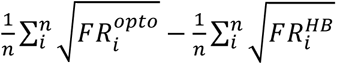, where 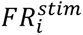 is the mean firing rate of unit *i* for that stimulus across repetitions, to estimate the difference in firing rates on average between the two stimulus conditions.

### Principal Components Projections

We project the simultaneously recorded neural data into a low dimensional projection as previously^50^. Briefly, we first compute the binned spike counts for each unit in 5ms bins in the last 5 minutes of the baseline recording. The binned spike counts were smoothed with a gaussian kernel (*σ* = 10*ms*) and square root transformed before applying PCA. In downstream analyses, we truncate the projection to a specific number of leading dimensions. We compute instantaneous trajectory speed as the Euclidean distance between two subsequent points (*dt* = 5*ms*) in that truncated projection in time. We then compute the post-stimulus point as the point in the latent space at a given time after the offset of the stimulus, for each stimulus repetition *i*. To compute dispersion (i.e., how far apart are the post-stimulus points), we create a matrix of all stimulus points (# projection dimensions X # stimulus repetitions), compute its covariance matrix Σ, then compute *Tr*(Σ). We compute the distance to inspiration-off by first computing the mean point in the latent space where respiratory phase is immediately after the offset of inspiration 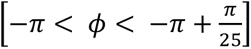 in the last 100 second window of the baseline recording (i.e., no stimulus). We compute the mean Euclidean distance to all post stimulus points as the mean distance to I-off. All computations are performed in the truncated low-dimensional space indicated. Random shuffled controls were computed by choosing 100 uniformly distributed random points from the last 100 seconds of the baseline window to be the stimulus offset times. Because there is a delay between closing of the valve and buildup of pressure in the lung, we consider the post-stim time to be 2.5 seconds after the valve closing.

### Dynamical modeling

To account for temporal structure when fitting a latent representation of the neural population activity, we fit recurrent switching linear dynamical systems models (rSLDS)^64^ as in^50^. Briefly, the rSLDS model fits a low dimensional continuous latent state *x*_*t*_ where *x*_*t*_ ∈ *R*^*d*^. The rSLDS also partitions the space such that the state of the model can be in any of *k* discrete states *z*_*k*_, and the discrete state *z*_*t*_ is a function of the continuous state *x*_*t*_. These partitions are each governed by separate linear dynamical systems subject to external inputs *u*_*t*_ such that the continuous state follows:

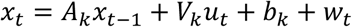

Where the fitted parameters are: *A*_*k*_, the dynamics matrix, *V*_*k*_ the control matrix applied to the input *u*_*t*_, and *b*_*k*_ the offset vector, all associated with discrete state *k*. *w*_*t*_ is a noise term which we set to zero during simulation due to the observed stereotypy of the neural activity. We choose *k* = 2, *d* = 2, *dt* = 10*ms* as we previously report this simple system to be sufficient to model these populations^50^. *u*_*t*_ is one dimensional and binary; it takes a value of 1 when the optogenetic laser is on, 0 otherwise. Thus *V*_*k*_ is a *d* dimensional vector. For analyses in Figure 7 where we analyze the learned stimulus field *V*_*k*_, we fit jointly {*A*_*k*_, *b*_*k*_, *V*_*k*_} during the last 5 minutes of the baseline period and the brief pulse stimulations only (50 repetitions each of 10ms and 50ms optogenetic stimuli). In analyses in Figure 8 where we impose arbitrary *V*_*k*_, we fit only {*A*_*k*_, *b*_*k*_} during the last 5 minutes of the baseline period (i.e., fit without optogenetic stimulus). While *u*_*t*_ is {0,1} during fitting, the term *V*_*k*_*u*_*t*_, the effective stimulus field during the *k*_*th*_ discrete state is uniform over the *k*_*th*_ partition, and differs in magnitude and direction for each discrete state. We restrict all subsequent analyses only for recordings that exhibited limit cycle dynamics (n=16 for joint fitting of all parameters, n=18 for fitting baseline only). Importantly, we showed previously that when limit cycle dynamics were observed, the two discrete states *z*_*k*_ corresponded to inspiration and expiration. We compute the stimulus strength for a given phase as the norm of the real components of each *V*_*k*_. Next we compute the eigenvectors of the dynamics matrices *A*_*k*_ and determine slow and fast eigen vectors for each discrete state by the magnitude of the eigenvalue. To determine how aligned the dynamics matrices *A*_*k*_ were with the stimulus vector *V*_*k*_, we compute the real component of the dot product between the normed eigenvectors and the normed stimulus vectors for each discrete state. Of particular importance was the slow expiratory eigenvector as it corresponded to the steady progression during expiration towards the next breath.

After fitting {*A*_*k*_, *b*_*k*_ } and optionally *V*_*k*_, we could simulate latent states *x*^ by initializing at *x*^ = (0,0) and following the latent state forward in time as it reached the limit cycle attractor according to the state update rule. Then, simulated integrated diaphragm traces were generated by first fitting a nonlinear support vector regression (SVR) with the fitted 2D latent state *x* as input and the observed integrated diaphragm trace as the target followed by applying that SVR to *x*^. We simulated optogenetic stimulation by multiplying *V*_*k*_ (either fitted from data or arbitrarily applied) by a chosen scaler multiplier that represented the stimulus amplitude. We performed simulated phasic triggered simulations by setting a threshold on the simulated integrated diaphragm above (or below) which we multiply *V*_*k*_ by some constant. We further controlled whether phasic stimulation occurred during the first (i.e., rising) or latter (i.e., falling) part of the inspiration, given the sub-phase specificity of NTS^GABA^ effects on breathing.

We manually examined a variety of arbitrary stimulus fields *V* (e.g, uniform across all directions, piecewise uniform, sinks, sources). We chose three simple stimulus fields that qualitatively matched the effects of the optogenetic stimulation observed in experiment: VRC^VGLUT2^-like, VRC^GABA^-like, NTS^GABA^-like. VRC^VGLUT2^-like was uniform in the direction of the slow expiratory eigenvector (constructive) with a sigmoidal decay in amplitude at the boundary of the inspiratory/expiratory partition. VRC ^GABA^-like was the mirror of VRC^VGLUT2^-like (opposed to the slow expiratory eigenvector). NTS^GABA^-like was also opposed to the expiratory eigenvector but constant amplitude over the entire space. The choice of a sigmoidal decay and the parameter of the decay was chosen manually that qualitatively matched observed data. To make stimulus amplitudes approximately comparable across recordings, stimulus amplitudes were scaled by the mean speed of the latent trajectory. Note that the amplitude scaling in the different fitting procedures (with or without including stimulus observations) are not comparable due to the explicit fitting of *V*_*k*_ in the former case. In some analyses, outlier data points were omitted after visual inspection showed that, for some recordings, the applied stimulus showed unexpected divergence or convergence to fixed points. These points are shown in figures, but excluded from statistical analyses.

### Statistics

Statistical analyses were computed using custom python software using statsmodels and pingouin. Analysis software is available at https://github.com/nbush257/VLAD.

### Software and data availability

All analysis software and documentation will be available on GitHub at https://github.com/nbush257/inspiration-off-attractor. Preprocessing software is available at https://github.com/nbush257/cibrrig, and experimental control software is available at https://github.com/nbush257/pyexpcontrol. Extracted data (spike data, physiology, events) in ONE format will be hosted upon publication. Raw electrophysiological data are available upon reasonable request.

## Discussion

### Overview

Here we provide a data-driven, low dimensional dynamical systems framework for flexible respiratory control *in vivo* informed by high-spatiotemporal resolution neural recordings and causal manipulations of identified respiratory control circuit sub-populations. These results further the inspiration-off switch mechanism as a central respiratory pacing mechanism *in vivo* as originally proposed by systems-level approaches. In these original investigations of the inspiratory-off switch mechanism^53,56,72,73^, the primary read-out has been gross physiological activity (i.e., EMG, airflow, nerve recordings, few single units) which represents the integrated output of a complex, heterogeneous, and spatially distributed neuronal network. These approaches allowed limited insights into the underlying neural mechanisms, and much of the support for these models was primarily inferred from computational biophysical simulations^74–76^. Here, we leverage comprehensive, high-resolution electrophysiological recordings with Neuropixels and the optogenetic manipulation of genetically defined neuronal subtype to unveil the neuronal network dynamics underlying flexible respiratory control.

This “inspiratory-off attractor” model provides further evidence for, and follows in spirit from, the classical inspiratory off-switch model in that timed phasic inhibition from mechano-sensory feedback resets the respiratory cycle to set breathing frequencies. It further augments the model in that it generalizes to predict bidirectional respiratory control from inhibitory populations, to account for manipulations of excitatory populations, and to predict the activity of the entirety of VRC neural populations. This is especially powerful as this generalization held even for predictive models fit without observations of stimulations.

Although, the simultaneous high-resolution recordings from large populations of distinct, genetically defined neuronal subtypes and their phase-specific selective manipulations constitute a fundamental departure from the previous, traditional in vivo studies, the present study clearly builds on important insights that were gained by many of these original studies. Here we show that breathing phase can be both advanced and delayed as a result of well-timed activity of inhibitory neurons in the NTS. Our experiments build on prior investigations of inhibitory cells in this region of the NTS, “pump cells”, that have been shown to receive direct innervation from slowly adapting lung mechanoreceptors^61,77,78^. These populations have been well implicated as respiratory interrupting, direct bidirectional control via optogenetics has not yet been shown. Indeed, because respiratory rhythmic activity can be maintained *in vitro* without this off-switch mechanism, there has been considerable uncertainty and confusion in the field surrounding the role and importance of the off-switch mechanism as proposed for the *in vivo* network. Some work speculated that the burst-termination as seen in vitro may reflect a “fail-safe” mode that could play a role in gasping but not eupneic breathing^79^.

It is known that in the absence of vagal inputs, respiration slows and diaphragmatic amplitude increases dramatically^67^. Normal rates and amplitudes can be recovered by direct stimulation of the remaining vagal nerves^72^ or by direct stimulation of inhibitory neurons of the preBötC^49^. Whether activation of the NTS^GABA^ populations in conditions of vagotomy are sufficient to recover qualitatively normal respiration is yet unknown, but an important next step. We speculate that the diverse molecular and anatomical targets of the vagi into the NTS make this plausible, but unlikely to be the sole circuit involved. For example our study did not address the role of the Kolliker-Fuse as an important contributor to the inspiratory off-switch. There are likely multiple pathways that can evoke identical, subtly different, or entirely different mechanisms that result in a similar physiological output, emphasizing the need for detailed and comprehensive neural recordings to disentangle. Indeed, our data here show similarity between physiological responses NTS^GABA^ stim and lung-inflation but subtly different effects on the VRC (namely, a subpopulation of inspiratory VRC neurons that are unaffected in lung-inflation). We expect that emerging techniques to target sub-populations based on more granular cell marker will provide critical insights into the mechanisms that govern the generation of breathing in the largely intact animal^80–82^.

### Subphase specificity of respiratory modulation by different sources of VRC inhibition

Interestingly, we find that activating NTS^GABA^ populations plays a significant role in inspiratory rhythmogenesis *in vivo* by specifically advancing the respiratory phase during the rising half of the inspiration while delaying phase in the second half of inspiration. This contrasts with direct stimulation of VRC^GABA^ populations which advances phase throughout inspiration. We interpret this as having two critical implications. Firstly, this agrees with the accepted role of the NTS^GABA^ population as relaying lung volume for the purpose of pacing via feedback inhibition and volutrauma protection. But, NTS^GABA^ neurons also amplified the re-afferent representation of lung volume, and the inspiration is terminated once this lung state representation reaches a threshold. Subsequently, amplification of this signal during early inspiration will pre-emptively trigger that reset. In contrast, amplification of the re-afferent signal in the latter half of inspiration will relay a lung-state representation that is larger than normal (as in a sigh), or in the inter-breath interval, a lung that is overfull (as in the Hering-Breuer reflex) and delay respiratory phase. Secondly, the fact that activating re-afferent circuitry during the latter half of inspiration delays respiration implies a role for inhibitory signaling in the VRC as rhythm advancing separate from mechanosensory feedback. Taken with the observation that latency to next breath after stimulation (i.e. rebound latency) is shorter for direct VRC^GABA^ stimulation than for NTS^GABA^, we conclude that inhibitory signaling within the VRC concurrently prepares for an inspiration while also opposing its execution.

The dynamical systems modeling results adds further evidence for this functional separability of inhibition. We showed that NTS^GABA^ like effects could be recreated by opposing the phase advancing dynamics (Figure 8) while VRC^GABA^ like effects were matched by opposing phase advance only during inspiratory dynamics. Consequently, during VRC^GABA^-like stimulation, the model systems are held at the edge of the transition into inspiration. This implies that VRC^GABA^ stimulation, but not NTS^GABA^ stimulation, prepares the dynamics to transition into inspiration. Moreover, VRC^GABA^ stimulation concurrently reduced the activity of expiratory unit activity (Figure 3F), while NTS^GABA^ did not (Figure 3F). It is important to note that these data do not reveal if it is specifically inhibitory, expiratory neurons that are inhibited here, we have previously shown that expiratory neurons are more likely inhibitory *in vivo*^50^. Our data are consistent with the notion that the VRC^GABA^ stimulation disinhibits the phase advancing components of the respiratory network that is absent during NTS^GABA^ stimulations. The separability of these inspiratory phase promoting and phase advancing components of inhibitory signaling suggests that purely excitatory mechanisms of rhythm generation, while well established as being sufficient for *in vitro* rhythms^41,43,83^, are likely less important during eupneic breathing *in vivo*. *In vitro* respiratory rhythms rely on recurrent excitability within the preBötC and are terminated by intrinsic cellular mechanisms and local interactions^38^. Our results offer an integrated view of the rhythmogenic process *in vivo* which relies on the dynamic, well-timed activation of inhibitory neurons to coordinate inspiratory motor commands with proprioceptive inputs derived from lung-afferents via the NTS. Further investigations will be important to unravel exactly what sub-circuits and cell-types might be involved in the phase promoting aspects of VRC inhibition.

In addition, we found that for our generative dynamical model to consistently recreate respiratory rate increases during inspiratory stimuli for the NTS^GABA^-like stimuli, we had to restrict stimulation to the rising phase of the diaphragm (Supplemental Figure 6C). This explains the difference in these results and recent reports in which inspiration timed NTS^GABA^ stimulations resulted in respiratory slowing^58^. In those results, the phasic stimulations were long enough to potentially continue during expiration.

### Antagonistic control of inspiratory transitions by expiratory/inspiratory balance

The Hering-Breuer reflex has been described for over a century^63^, and the responses of single neurons of various subregions of the VRC have been well characterized^51,57,62,84–89^. However, the coordinated changes in activity that arise in these distributed neural populations of the VRC during specific activation of transcriptionally defined respiratory populations has not been investigated. Interestingly, we observe that during lung inflation, expiratory neural activity remains high until a breath is triggered (Figure 5G, K); contrasting the observation that expiratory neural activity decays in advance of the increase in inspiratory populations (Figure 5G). The lung inflation keeps expiratory tone high, which in turn putatively inhibits inspiratory tone. Simultaneously, inspiratory tone increases over time, likely due to a combination of intrinsic excitability^90,91^ and tonic excitatory input^92^. Whether and when the inspiratory tone overcomes this balance will likely be specific to the sensory input and internal state of the animal. Lung inflation opposes inspiratory tone and recruits expiratory neurons, and the inspiratory drive to overcome that expiratory “barrier” is increased. This “stiffening” of the transition to inspiration is supported by the observation of a coordinated increase in both inspiratory and expiratory firing rates just before inspiration (Figure 5K), with the absence of a change in the ratios of firing (Figure 5M). Further supporting this is the change in inspiratory dynamics during lung inflation. Inspiratory neural activity is higher (Supplemental Figure 5F), inspirations are shorter (Figure 5E), and neural trajectories during inspiration are faster (Supplemental Figure 5G). In other systems (e.g., arm movement^93^ and fish locomotion^94^), a similar mechanism of antagonistic control is a common strategy for fine-tuned control. Balance in two opposing forces results in stability, while imbalance moves the state with more precision than without antagonistic control.

It is important to note the (expected) lack of a perfect match between neural activity during Hering-Breuer reflex responses and stimulation of either NTS^GABA^ or VRC^GABA^ populations (Figure 6H). This indicates that the optogenetic stimulus cannot completely mimic the lung inflation stimulus at the detailed level of neural populations. The lung-inflation stimulation will specifically activate an endogenous inflation protection circuit, which will include activation of rapidly adapting stretch receptors (RARs)^95,96^ and the pons^73^. Conversely, the optogenetic stimulations will likely activate off-target circuits which may be involved in other aspects of respiratory control, or other, non-respiratory functions^97^. The strong match in the breathing data for both stimulus types without the perfect match of the neural data is a warning that behavioral/gross physiological readouts are not complete, and limits how much certainty can be said without comprehensive examination of the neural activity.

### Contrasting dynamic motifs to modulate neural rhythms

Our data show that stimulation of VRC^VGLUT2^, VRC^GABA,^ or NTS^GABA^ populations can differentially increase respiratory rate if done with precise timing. By interrogating latent population dynamics, we uncover that the neural mechanism by which this respiratory rate increase occurs is strikingly different. The inspiration-off target is a critical feature in this dynamic landscape. Neural trajectories are attracted to, and dwell in, this region. Activation of VRC^VGLUT2^ populations speeds up neural trajectories by pushing them away from the inspiratory off target (i.e. destabilizing); conversely, activation of either inhibitory population temporarily increases the attractiveness of that region (i.e., stabilizing). In this way, VRC^VGLUT2^ populations increase respiratory rate by increasing trajectory speeds, most strongly in between inspirations, while inhibitory stimulations “short cut” the latent trajectory. This perturbation of the dynamic landscape is further supported when the latent dynamics are modeled explicitly and the impact of driving input or constituent subcircuits can be mapped directly onto vector fields with those behaviors that act on the neural dynamics.

Importantly, in the transgenic (VRC^GABA^ or VRC^VGLUT2^) mice, we stimulate not only local VRC neurons, but also glutamatergic or GABAergic inputs, and therefore are activating broadband excitatory and inhibitory inputs to the region. We show here that lung stretch inputs are sufficient to drive the network back to the inspiration off attractor and likely play a primary role in shaping the normal eupneic dynamics. However, further experiments are required to delineate circuit, region, or input cell type specific roles in sculpting or modulating those dynamics. In particular, we hypothesize that glutamatergic inputs from NMB^+^ RTN neurons may play a primary role in destabilizing the inspiratory off attractor, particularly that inhibiting and removing these populations drastically reduces the regularity and rate of the inspiratory rhythm *in vivo*^12^.

Notably, VRC^GABA^ stimulation more quickly and directly drove the dynamics to the inspiration off target than did NTS^GABA^ stimulations. This agrees with our targeting of the NTS^GABA^ neurons that are presumptive secondary afferents receiving input from the SARs and affect breathing potentially via a multi-synaptic pathway^57,98,99^. Recent compelling work has implicated local inhibition of NTS^Phox2B^ neurons^58^, but they do not rule out the function of additional collateral projections. Other possible mechanisms for these slower perturbation responses may include slower intrinsic dynamics of NTS^GABA^ cells, smaller post-synaptic effects, or lower levels of transfection in viral approaches compared to transgenic.

## Conclusion

The conceptual model for respiratory control as presented here provides a differentiated view of the rhythmogenic process in which inhibitory and excitatory mechanisms play opposing roles in the coordination of the respiratory cycle via dynamic control of the stability of an inspiratory off attractor. Further, this control is in large part dictated by mechanoreceptive inputs from lung afferents. However, our study also raises important questions that remain unanswered. First, how is the lung stretch set point determined? What exogenous or endogenous subcircuits within the medulla, pons and other areas of the brain can manipulate that set point? What are the contributions of various inspiration promoting mechanisms to the destabilization of the inspiratory-off fixed point? We speculate that excitatory chemo-sensitive inputs from the carotid bodies and the RTN/pFRG are also critically involved. Lastly, how do other inspiration terminating mechanisms (e.g., those triggered by irritants or e.g. during speech/vocalization ^13,100 15^) or slowing mechanisms^19^ affect VRC latent dynamics?

Together these data emphasize the role of inhibitory re-afferent feedback in shaping the normal ongoing dynamics of diverse respiratory motor control populations that govern breathing. Viewing the activity of neural populations in a low-dimensional dynamic space allows us to decompose the impact of manipulating cell-type specific subcircuits and inputs into the respiratory control circuitry. This has been used to great effect in other motor and behavioral settings^101,102^, offering an exciting and powerful way to dissect neural circuits in exquisite detail beyond observing motor output (e.g., breathing behavior). When combined with perturbation of intersectionally defined cell-type and projection specific populations^45,46,103^, we can begin to disambiguate the functional role of neural populations that may have overlapping or indistinguishable impact on respiratory output.

## Acknowledgements

We thank the National Institutes of Health for funding: NIH R01 HL144801, R01 HL126523, and R01 HL151389 to J.M.R, NIH F32HL159904 to N.E.B. We thank the Behavioral Phenotyping Core (RRID:SCR_026371) at Seattle Children’s Research Institute for its assistance with the completion of neural recordings. We thank Research Scientific Computing at Seattle Children’s Research Institute for providing HPC resources that have contributed to the research results reported within this paper.

## Declaration of generative AI and AI-assisted technologies in the manuscript preparation process

During the preparation of this work the author(s) used GitHub Copilot in order to aid writing of analysis software. After using this tool/service, the author(s) reviewed and edited the content as needed and take(s) full responsibility for the content of the published article.

## Supplemental Figures

**Supplemental Figure 1:**
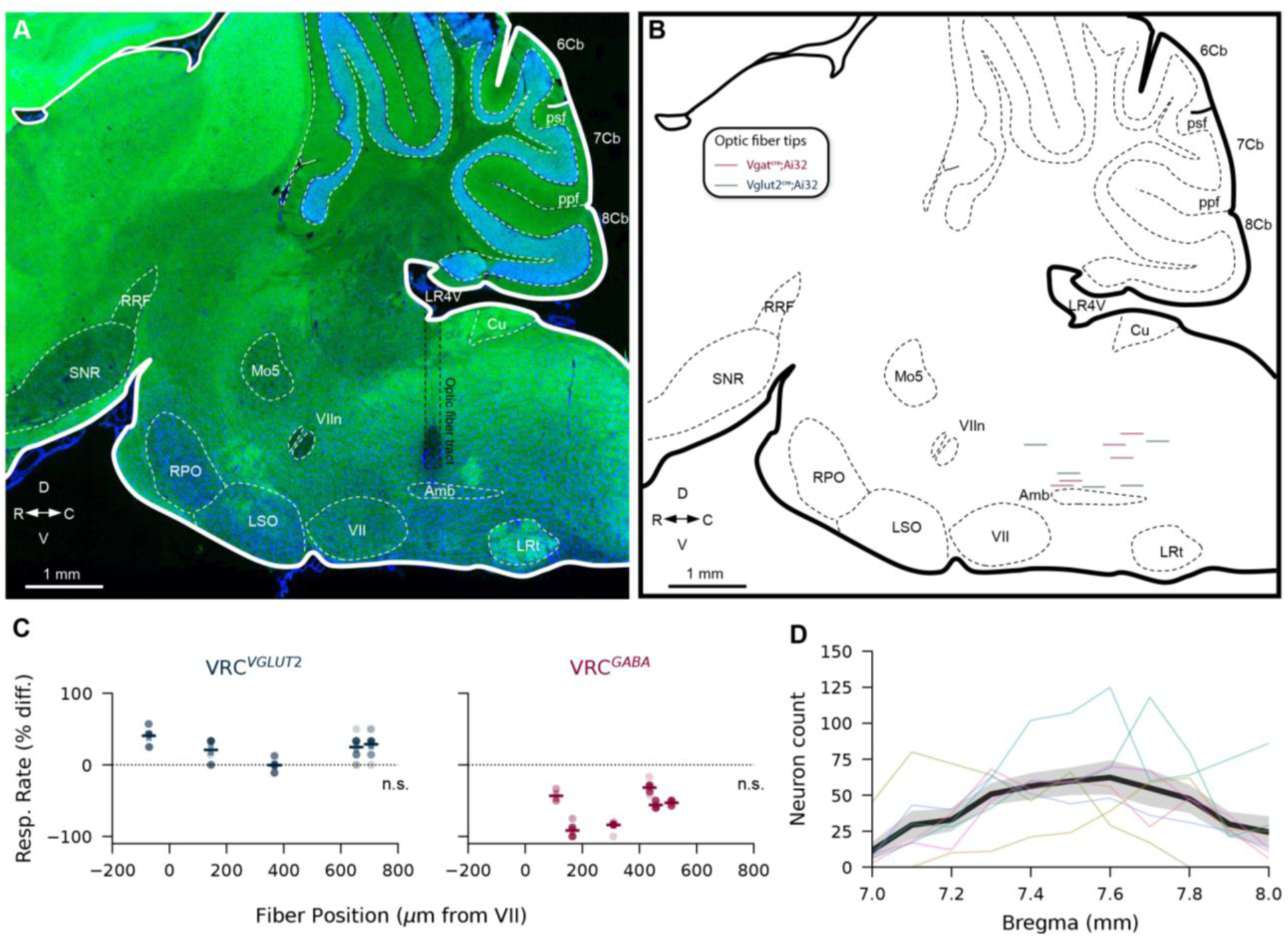
(A) Example histology and summary (B) of optic fiber locations for VRC^GABA^ (dark red) and VRC^VGLUT2^ (dark blue). (D) Change in respiratory rate as a function of fiber placement for 2s hold stimulations. Horizontal lines are mean of each animal, vertical lines are S.E.M., dots are individual stimulations. (Mixed linear model VRC^VGLUT2^ z=-0.430 coeff 95%CI= [-0.061,0.039], p=0.667, n=5, VRC^VGAT^ z=0.759 coeff 95%CI= [-0.079,0.179], p=0.448, n=6) (D) number of cells transfected with ChRmine in NTS^GABA^ animals as a function of bregma. Black line is average, shaded region is S.E.M, colored lines are individual animals (n=7). Abbreviations: Amb, ambiguus nu; 6Cb, 6th cerebellar lobule; 7Cb, 7th cerebellar lobule; 8Cb, 8th cerebellar lobule; Cu, cuneate nu; LR4V, lateral recess of 4^th^ ventricle; LRt, lateral reticular nu; LSO, lateral superior olive; Mo5, motor trigeminal nu; psf, posterior superior fissure; ppf, prepyramidal fissure; RPO, rostral periolivary region; RRF, retrorubral field; SNR, substantia nigra reticular; VII, facial nu; VIIn, facial nerve.

**Supplemental Figure 2:**
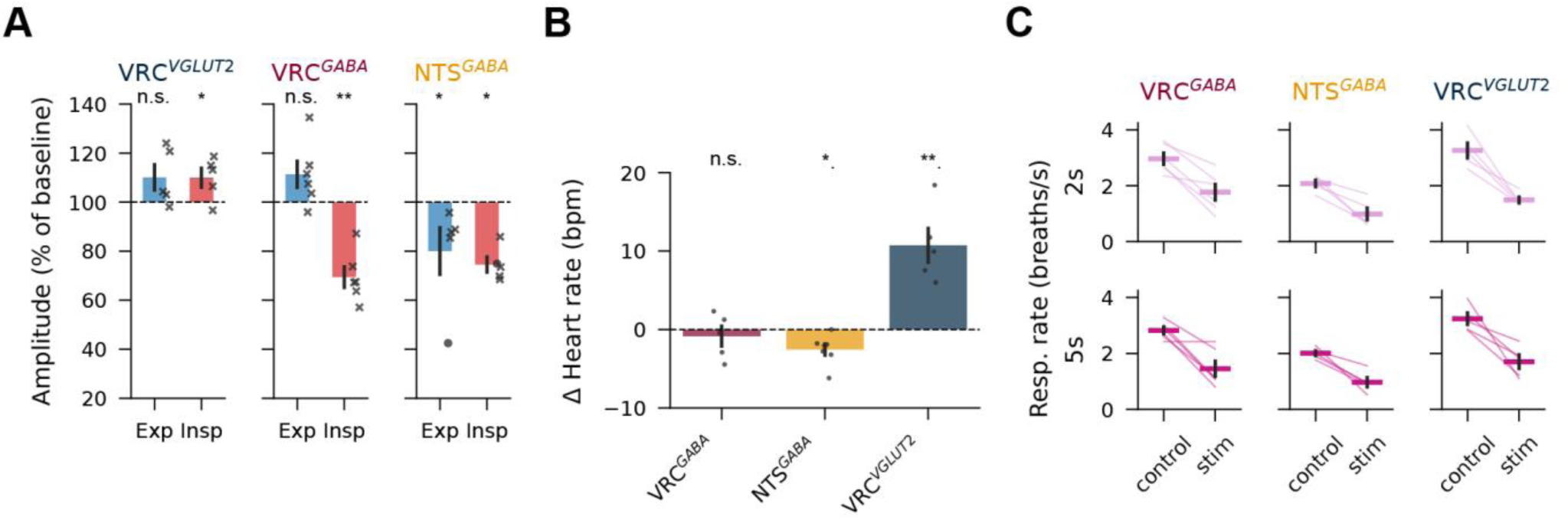
Physiological effects of optogenetic and physiological stimulations. (A) Change in diaphragm EMG amplitude (as % of baseline) for phasic-triggered optogenetic stimulations, stratified by targeted population. (One sample t-test VRC^VGLUT2^ n=5, p_insp_=0.04, p_exp_=0.17; VRC^GABA^ n=6, p_insp_=0.007, p_exp_=0.09; NTS^GABA^ n=5, p_insp_=0.028, p_exp_=0.046). (B) Change in heart rate (beats per minute) during 2s hold optogenetic stimulations, stratified by targeted genotype (Two way paired t-test pre stimulus compared to during stimulus VRC^VGLUT2^ n=5, p_insp_=0.007; VRC^GABA^ n=5, p_insp_=0.054; NTS^GABA^ n=7, p=0.01). (C) Respiratory rate before (control) and during Hering-Breuer stimulation (lung inflation) held for 2s or 5s (row) and stratified by optogenetic target (columns). Three-way ANOVA (stimulus,target,duration). significant main effect of stimulus p_stimulus_=2.19E-15,F_stimulus_=124.1 and target p_target_=2.2E-7, F_target_=20.8, but not duration p_duration_=0.55,F_target_0.35; no significant interaction effects see supplemental table 1) Bars are mean +/- S.E.M.

**Supplemental Table 1:** Hering-Breuer ANOVA table.

|  | sum_sq | df | F | PR(>F) |
| --- | --- | --- | --- | --- |
| *C(target)* | 9.5287 | 2 | 20.87395 | 2.21E-07 |
| *C(duration)* | 0.079806 | 1 | 0.349653 | 0.556871 |
| *C(stimulus)* | 28.32901 | 1 | 124.1173 | 2.19E-15 |
| *C(target):C(duration)* | 0.295684 | 2 | 0.647736 | 0.527398 |
| *C(target):C(stimulus)* | 0.88583 | 2 | 1.940535 | 0.153888 |
| *C(duration):C(stimulus)* | 0.002756 | 1 | 0.012076 | 0.912919 |
| *C(target):C(duration):C(stimulus)* | 0.11048 | 2 | 0.242023 | 0.785918 |
| *Residual* | 11.86868 | 52 |  |  |

**Supplemental Figure 3:**
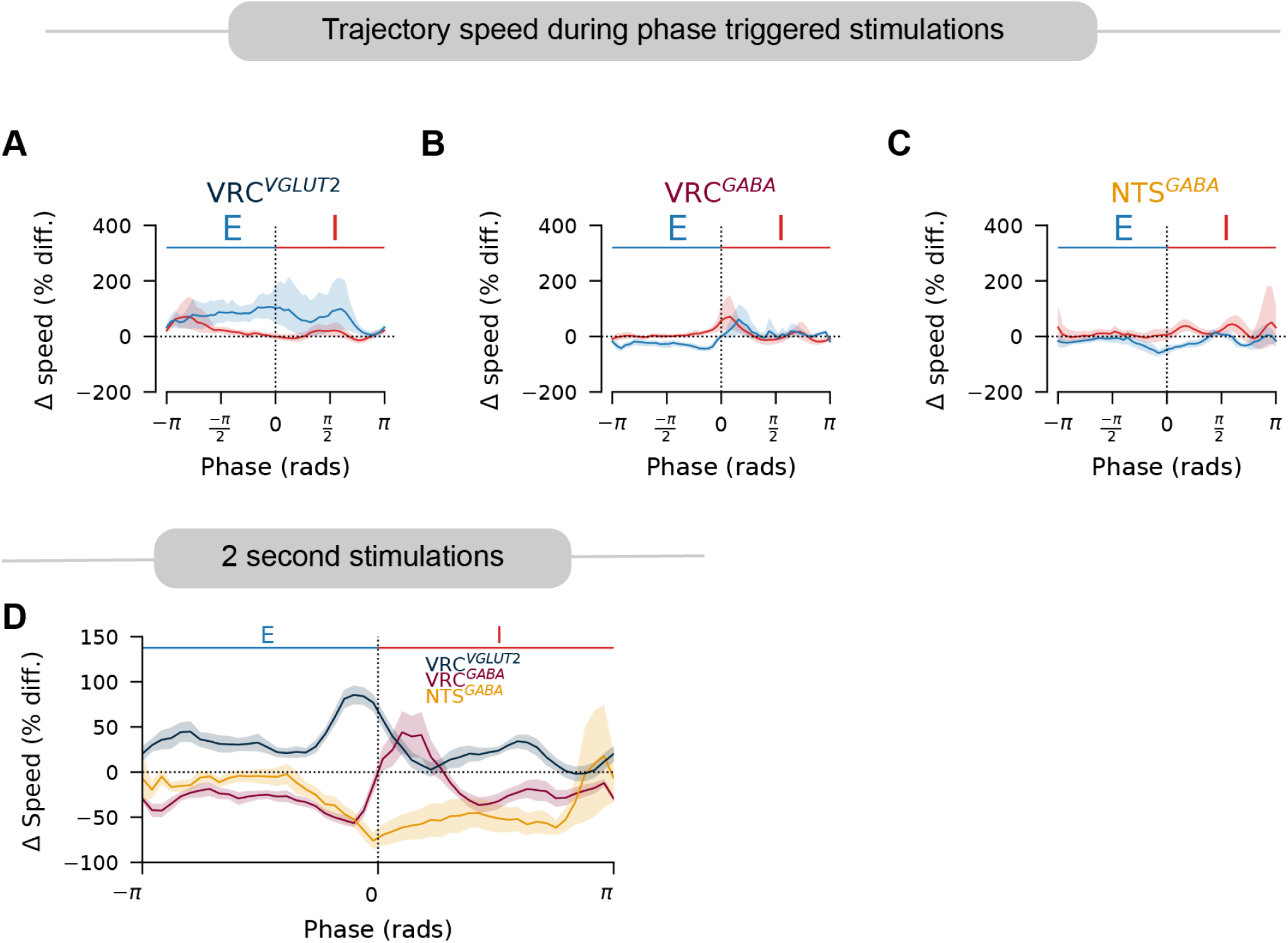
Change in trajectory speed during optogenetic stimulation of different target populations. (A-C) Phase averaged trajectory speed during phasic-triggered stimulations. Shaded regions are mean +/- 95% C.I. VRC^VGLUT2^ n=5, VRC^GABA^ n=6, NTS^GABA^ n = 4. (D) Phase averaged trajectory speeds for 2s hold stimulations. Shaded regions are mean +/- S.E.M. VRC^VGLUT2^ n=5, VRC^GABA^ n=6, NTS^GABA^ n = 5.

**Supplemental Figure 4:**
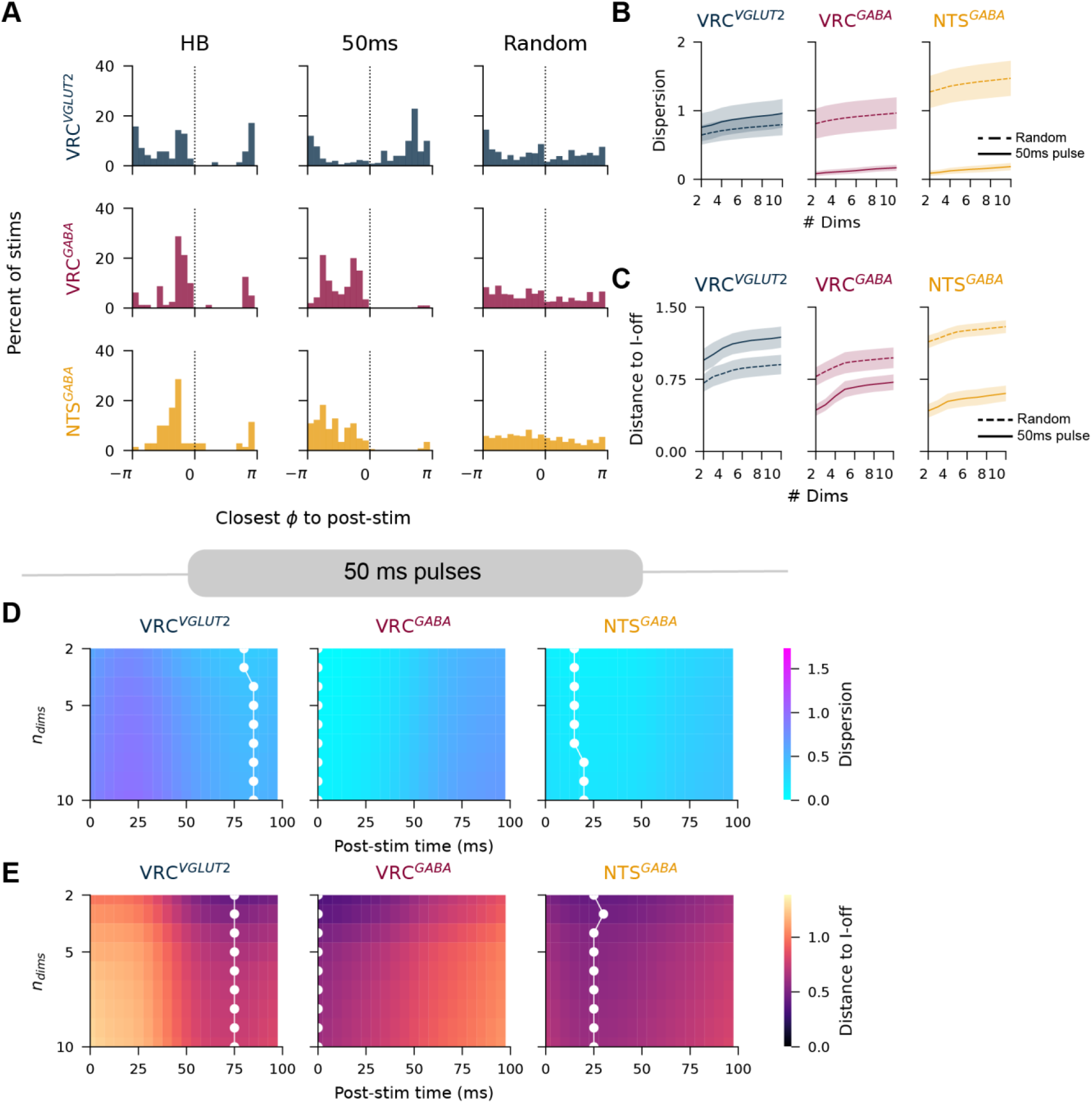
Convergence of latent trajectories after short optogenetic pulses. (A) Histogram of the values of phase from baseline average trajectory nearest to the average latent state during Hering-Breuer stimulation (HB), 25ms after brief 50ms optogenetic pulse (50ms) or random shuffled control points (random) aggregated across recordings. Rows stratified by optogenetically targeted population. (B) Dispersion and (C) distance to inspiration-off point of post-stimulus latent state 25ms after brief 50ms optogenetic pulse (solid line) or random baseline control points (dashed line) computed for increasing included dimensions of the latent state. Shaded regions are mean +/- S.E.M. (D) Dispersion and (E) Distance to inspiration off point as a function of both time after stimulation (x-axis) and number of dimensions included in latent state (y-axis) for 50ms optogenetic pulses, averaged across recordings. White dots indicate minimum along y-axis. VRC^VGLUT2^ n=5, VRC^GABA^ n=6, NTS^GABA^ n = 5.

**Supplemental Figure 5:**
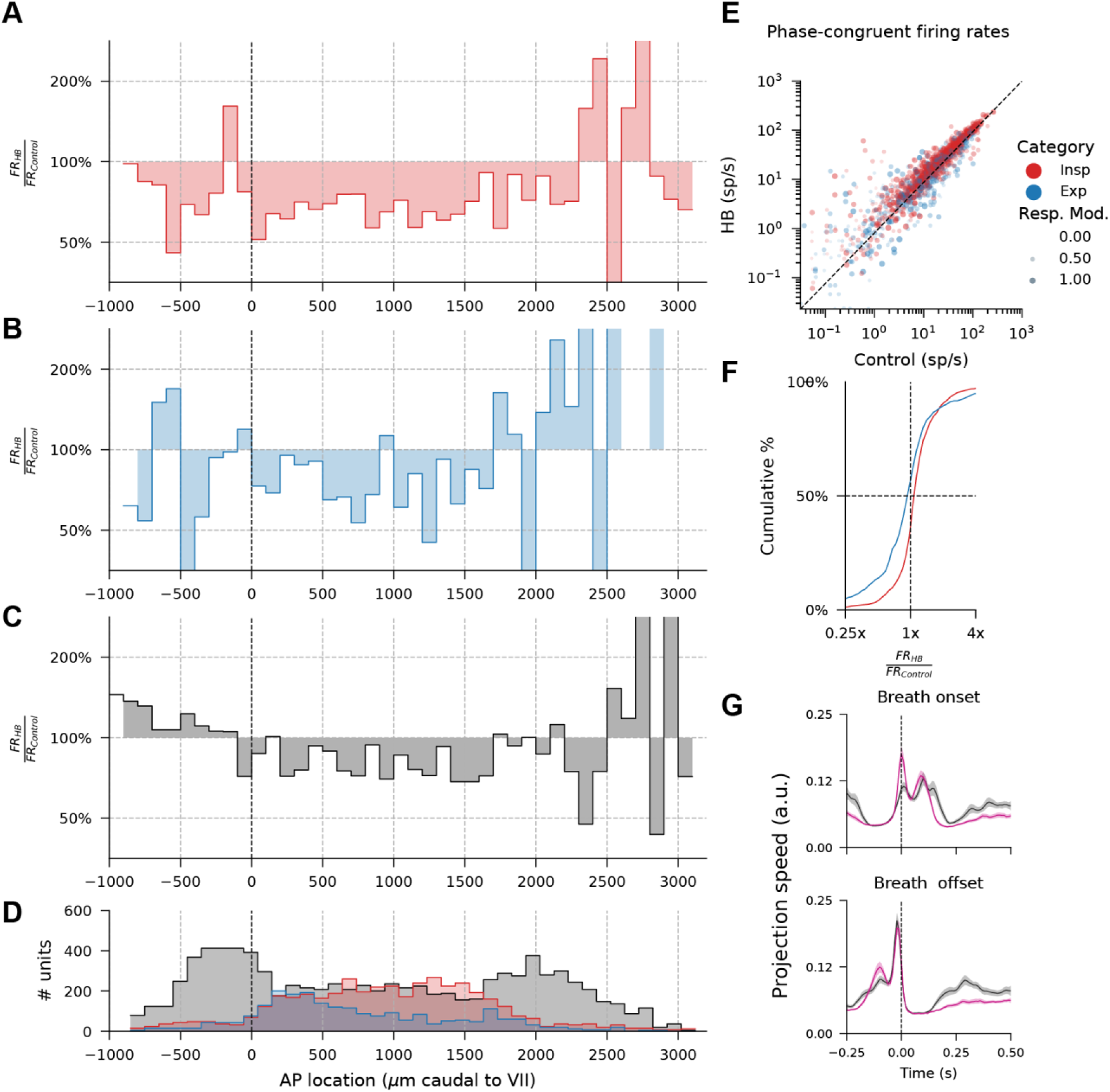
Changes in neural dynamics during Hering-Breuer stimulation. (A-C) Percent change in firing rate for (A) inspiratory (B) expiratory and (C) tonic neurons averaged in 100*μ*m bins as a function of inferred anterior posterior location relative to the caudal border of the facial nucleus (VII). (D) Number of units in each bin – fewer units near anterior/posterior extents contributes to nosy estimates of changes in firing rate. (E) Firing rate in baseline control (x-axis) vs Hering-Breuer stimulation(y-axis), but excluding time and spikes observed outside of preferred phase (i.e., inspiratory neuron firing rate is computed for spikes only in inspiration, and normalized only for inspiratory time). Each dot is a unit. (F) Cumulative distribution of firing rate ratios. Right shifted means neurons fire faster during Hering-Breuer stimulation (p_insp_=2.3E-39 W_insp_=150,296; p_exp_=0.039,W_exp_=74,336). (G) PCA trajectory speed aligned to (top) breath onset and (bottom) breath offset during baseline control (black) and break through breaths in Hering-Breuer stimulation (magenta). Shaded region is mean +/- S.E.M.

**Supplemental Figure 6:**
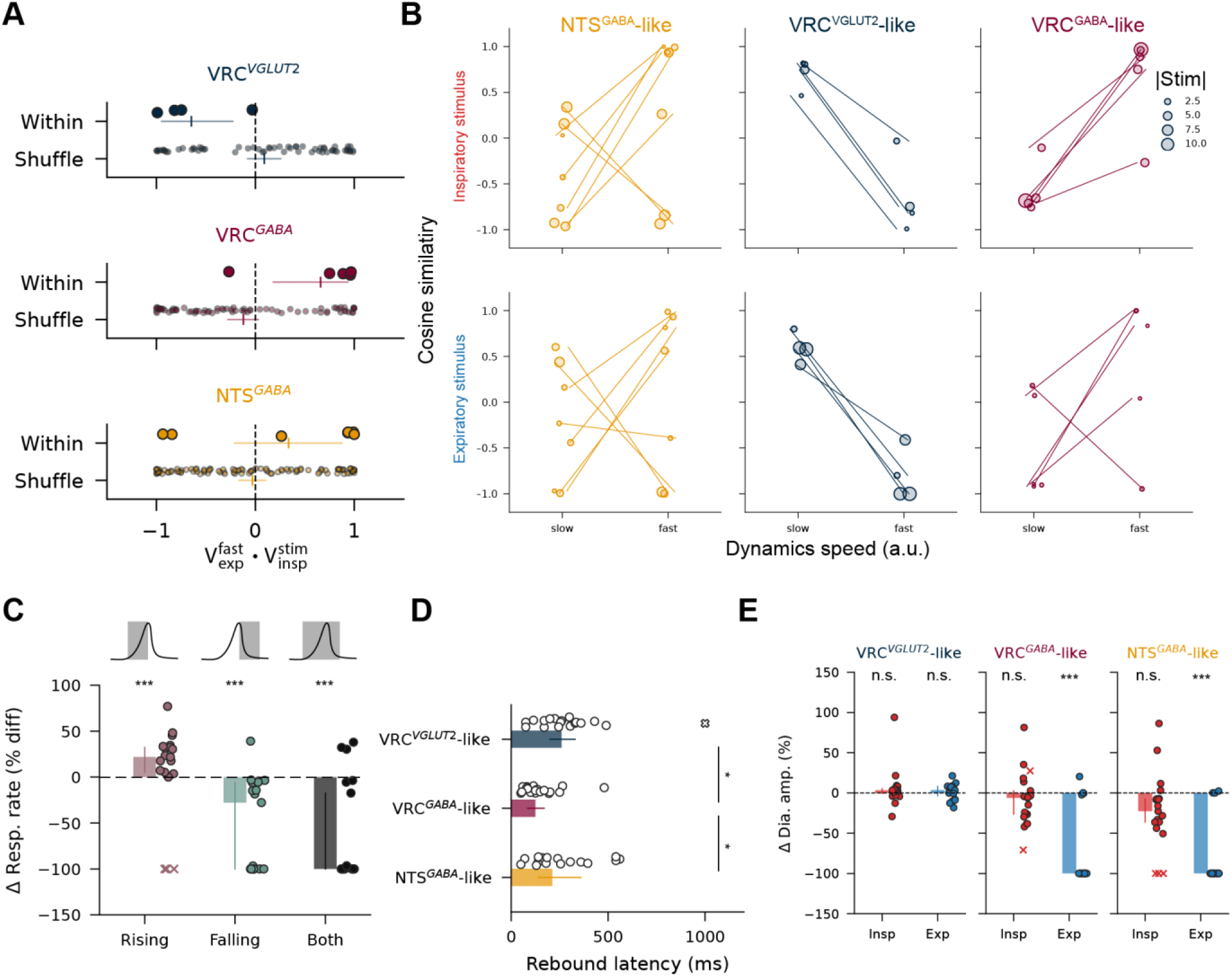
rSLDS simulations (A) Cosine similarity of learned inspiratory stimulus vectors with fast expiratory eigenvectors as in Figure 7E. (B) Cosine similarity of inspiratory (top) and expiratory (bottom) stimulus vectors with both slow and fast expiratory eigenvectors (includes data in Figure 7E and Supplemental Figure 5A). Dot size indicates norm of stimulus field. Lines indicate stimulus vectors from the same recording. Color signifies experimentally targeted populations. (C) Change in simulated respiratory rate during NTS^GABA^-like, inspiratory-triggered stimulations restricting stimulation to either the rising or falling period of the simulated diaphragm contraction, or to both periods. (one sample Wilcoxon signed rank test: p_rising_=6.5E-4, W_rising_=0, N_rising_=16;p_falling_=1.6E-4, W_falling_=10, N_falling_=19,p_both_=5.2E-4, W_both_=15, N_both_=19.) (D) Latency from offset of stimulus until next simulated breath onset for the three stimulus types (Friedman test dof=2, Q=17.1, p=1.8E-4, post-hoc test Holm-Bonferroni corrected *p<0.05). (E) Change in simulated diaphragm amplitude (as % of baseline) for phasic-triggered simulated stimulations, stratified by stimulus type. (One sample Wilcoxon, outliers as crosses defined from C, *** p<0.001, see supplemental table 2 for statistics). In all panels bars are median +/- 95% CI (outliers included), crosses indicate outliers.

**Supplemental Table 2:**
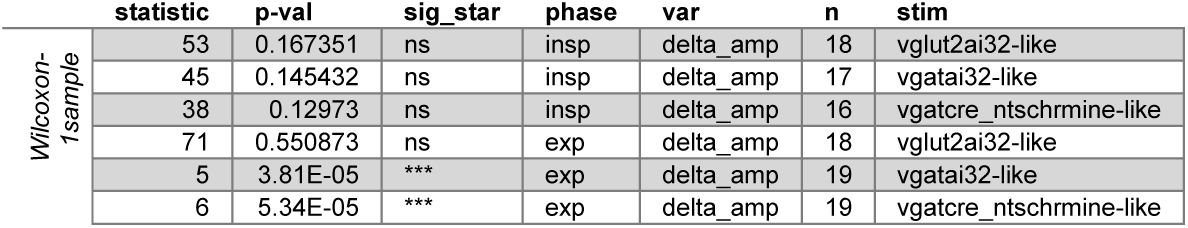
Phasic NTS^GABA^-like simulated stimulations statistics.

## Supplemental Videos

### Supplemental Videos 1-3

Effects of stimulation of VRC^GABA^, NTS^GABA^, and VRC^VGLUT2^ (Supplemental videos 1,2,3 respectively) on diaphragm activity (top, yellow), single unit activity (middle, white), and two dimensional PCA projection (bottom). Hold stimulations and inspiratory triggered stimulations are shown sequentially. Optogenetic activation is shown as a solid colored line above the diaphragm trace (color indicates applied laser color specific for appropriate expressed opsin). Light cyan traces in PCA projection are past, unshown activity, yellow traces are unstimulated trajectory, laser colored traces (blue or red) are trajectories during stimulation.

### Supplemental video 4

Effects of Hering-Breuer stimulation on diaphragm activity (top, yellow), single unit activity (middle, white), and two dimensional PCA projection (bottom) for the same recording as in Supplemental video 1. Positive pressure is shown as a solid magenta line above the diaphragm trace. Light cyan traces in PCA projection are past, unshown activity, yellow traces are unstimulated trajectory, magenta traces are trajectories during stimulation.

### Supplemental Videos 5,6

Simulated rSLDS model dynamics (two dimensional, two state) from example VRC^GABA^ and NTS^GABA^ recordings with fitted stimulus fields (Supplemental videos 5,6 respectively). Ramping stimulus amplitude and pulses of various durations are shown. Stimulus amplitude is shown top in yellow. Background color indicates vector field speed; apparent discontinuity in speed is the boundary between partitions of the latent state governed by the individual discrete states. Green dot and trail indicates simulated latent dynamical model with initial state (0,0).

